# Ovule siRNAs methylate protein-coding genes in *trans*

**DOI:** 10.1101/2021.06.10.447945

**Authors:** Diane Burgess, Hiu Tung Chow, Jeffrey W. Grover, Michael Freeling, Rebecca A. Mosher

## Abstract

24-nt small interfering siRNAs maintain asymmetric DNA methylation at thousands of euchromatic transposable elements in plant genomes in a process call RNA-directed DNA Methylation (RdDM). RdDM is dispensable for growth and development in Arabidopsis, but is required for reproduction in other plant species, such as *Brassica rapa.* 24-nt siRNAs are particularly abundant in maternal reproductive tissue, due largely to overwhelming expression from a small number of loci in the ovule and developing seed coat, termed siren loci. Recently it was shown that abundantly expressed 24-nt siRNAs produced in the tapetal tissue of anthers can methylate male meiocyte genes *in trans* (Long et al., 2021). Here we show that a similar process takes place in female tissue. siRNAs are produced from gene fragments embedded in some siren loci, and these siRNAs can trigger methylation in *trans* at related protein-coding genes. This *trans*-methylation is associated with silencing of some target genes and may be responsible for seed abortion in RdDM mutants. Furthermore, we demonstrate that a consensus sequence in at least two families of DNA transposons is associated with abundant siren expression, most likely through recruitment of the CLSY3 putative chromatin remodeller. This research describes a new mechanism whereby RdDM influences gene expression and sheds light on the role of RdDM during plant reproduction.

## INTRODUCTION

In plants, DNA methylation can be initiated *de novo* via RNA-directed DNA Methylation (RdDM), however how and where RdDM is initiated often remains mysterious (Matzke and Mosher, 2014). RdDM begins when RNA Polymerase IV (Pol IV) and RNA-DEPENDENT RNA POLYERMASE 2 (RDR2) produce short non-coding double-stranded transcripts that are processed by DICER-LIKE 3 (DCL3) into 24-nt short interfering (si)RNAs (Blevins et al., 2015; Li et al., 2015; Singh et al., 2019; Zhai et al., 2015). One strand of these siRNA duplexes is then loaded into ARGONAUTE 4 (AGO4) or its relatives (Havecker et al., 2010). The AGO/siRNA complex interacts with non-coding transcripts produced by RNA Pol V and recruits DOMAINS REARRANGED METHYLTRANSFERASE 2 (DRM2) to catalyze cytosine methylation (Böhmdorfer et al., 2014; Liu et al., 2018; Wierzbicki et al., 2009). RdDM primarily methylates small euchromatic transposable elements (TEs) and DNA present at the edges of larger TEs, particularly those closer to genes (Zemach et al., 2013). Despite its function in euchromatin, RdDM rarely functions at protein-coding genes (Matzke and Mosher, 2014).

Genomic regions are targeted for 24-nt siRNA production by members of the CLASSY (CLSY) family of putative chromatin-remodeling factors (Zhou et al., 2018). The four CLSY family members direct Pol IV to chromatin in a series of partially-redundant relationships. CLSY1 and CLSY2 function with SAWADEE HOMEODOMAIN HOMOLOGOUE1 (SHH1) and are responsible for nearly all 24-nt siRNA production in leaves (Law et al., 2013; Zhou et al., 2021). In parallel, CLSY3 and CLSY4 are responsible for Pol IV activity at a subset of loci in flowers, and CLSY3 is required for expression of abundantly-expressed 24-nt loci in ovules and meiocytes (Long et al., 2021; Zhou et al., 2021).

In most cases, 24-nt siRNAs target DNA methylation in *cis,* utilizing their perfect complementarity to maintain asymmetric CHH methylation after DNA replication (where H=A, T, or C). However, exogenous 24-nt siRNAs are capable of triggering DNA methylation and transcriptional silencing with up to 2 mismatches between the siRNA and target locus (Fei et al., 2021), indicating that Pol IV-derived siRNAs might also function in *trans.* It was recently demonstrated that 24-nt siRNAs produced in the tapetum move into meiocytes, where they *trans*-methylate TEs and protein-coding genes, triggering changes in gene expression during pollen development (Walker et al., 2018; Long et al., 2021). However, it is unclear whether *trans*-methylation of protein-coding genes is a pollen-specific phenomenon or occurs more broadly during plant development.

Mutation of the RdDM pathway has only subtle defects in Arabidopsis, while loss of RdDM in other species have dramatic impacts on reproduction (Chow et al., 2020). In *B. rapa,* which diverged from Arabidopsis approximately 14.5 million years ago (mya), loss of RdDM in the maternal sporophyte results in a high rate of seed abortion (Grover et al., 2018). Seed development is dependent on RdDM activity in the maternal sporophyte, suggesting that 24-nt siRNA production in diploid maternal tissues is critical for sustained development of the endosperm and/or embryo (Grover et al., 2018).

We previously characterized a set of loci with extremely high expression of 24-nt siRNAs in ovules (termed siren loci). Although siren loci comprise only 1-2% of all 24-nt dominant loci in *Brassica rapa (B. rapa)* ovules, they account for 90% of siRNA accumulation (Grover et al., 2020). In contrast, accumulation of siRNAs at these loci is negligible in leaves and anthers. Siren siRNA expression is dependent on Pol IV, RDR2, and CLSY3, and siren loci are heavily methylated in ovules and developing seed coats, demonstrating that siren siRNAs act in *cis* through canonical RdDM (Grover et al., 2020; Zhou et al., 2021). In endosperm, there is moderate accumulation of siren siRNAs, but with a striking maternal bias and a dependency on siren siRNA expression in maternal tissues (Grover et al., 2020), suggesting that these extremely abundant siRNAs might be transported into the endosperm from surrounding maternal tissues.

To better understand the origin and consequences of RdDM during *B. rapa* reproduction, we investigated siren loci and the resulting siren siRNAs. Here we show that *B. rapa* siren loci preferentially map to genic and intergenic regions and in most cases only overlap TEs at their edges. At least two TE families are associated with siren loci, suggesting that a specific region of these TEs may induce siren formation in the adjacent intergenic region. One-third of siren loci overlap genes or gene fragments, and in some cases, the regions in related *B. rapa* genes corresponding to these gene fragments are *trans*-methylated specifically in tissues where siren siRNAs are expressed. Transcript accumulation for some of these *trans*-methylated target genes is impacted by RdDM, suggesting that siren siRNAs regulate expression of protein-coding genes during reproduction.

## RESULTS

### Ovules and seed coats produce abundant non-TE 24-nt siRNAs

To better understand the relationship between TEs and siRNA production in reproductive tissues, we manually annotated 415 TE families in the *B. rapa* R-o-18 genome, including families that are lineage-specific and poorly-conserved. Approximately 40% of the *B. rapa* genome is composed of TE sequence compared to 21% for Arabidopsis (Wang et al., 2011). While 58% of leaf 24-nt siRNAs mapped to annotated TEs, only 13% of ovule 24-nt siRNAs mapped to TEs (**Figure 1A**). A similarly low percentage of seed coat 24-nt siRNAs mapped to TEs (12-13%), whereas the percentage was higher in endosperm (43%) and embryo (55%). Consistent with the low rate of TE-derived sequences, the percentage of 24-nt siRNAs that map to only one genomic position was higher in ovule and upper seed coat compared to leaf, endosperm, or embryo (**Figure 1B**).

**Figure 1.**
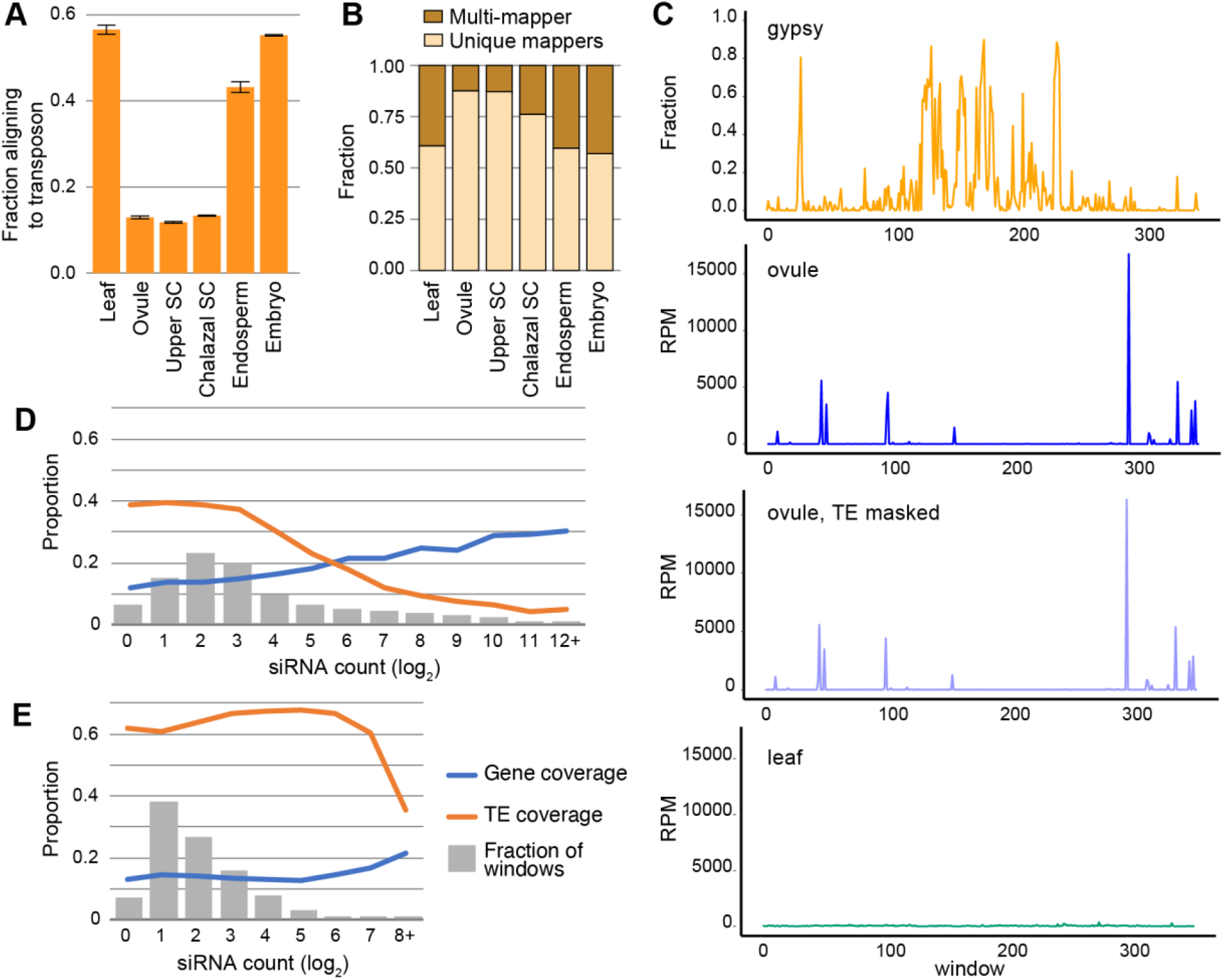
*B. rapa* ovule 24-nt siRNAs predominantly map to non-repetitive regions and siRNA hotspots. **(A)**The fraction of 24-nt siRNAs from various tissues and organs aligning to annotated TEs. Error bars depict the standard error of the mean. SC, 17 days post fertilization seed coat. **(B)**Fraction of 24-nt siRNAs from different tissues and organs mapping uniquely or to multiple genomic locations. **(C)**Distribution of *Gypsy* elements and 24-nt siRNAs across 100 kb windows of chromosome A01 of R-o-18, v2.3. Results for chromosomes A02-A10 can be viewed in Supplemental Figure 1. *Gypsy* retrotransposon data are plotted as the fraction coverage for each window. **(D, E)**Higher siRNA density in ovules correlates with higher gene coverage and lower TE coverage. 24-nt siRNA clusters from ovules or leaves were sub-divided into 24-nt sliding windows with a step size of 12-nt, and the average fraction covered by TEs or genes was plotted for each read count range (log2 scale). Bins with the highest-expressing windows were pooled until all bins contained at least 200 windows (12+ for ovules, 8+ for leaves). The fraction of windows in each bin is shown in gray bars.

Given the greater number of uniquely-aligning 24-nt siRNAs in ovules, we next asked whether these siRNAs differ in their chromosomal distribution in ovules and leaves. As a proxy for pericentromeric regions and large gene-poor heterochromatic regions, we used *Gypsy* retrotransposon coverage (Quesneville, 2020; Lim et al., 2005). In each chromosome, uniquely-mapping ovule 24-nt siRNAs occurred in discrete peaks that correspond to previously described siren loci (Grover et al., 2020) (**Figure 1C, Supplemental Figure 1A**). This pattern was highly reproducible (**Supplemental Figure 1B**) and persisted when reads were mapped to a TE-masked genome, but differs strikingly from uniquely-mapping leaf 24-nt reads, which align evenly but at a much lower density across chromosomes (**Figure 1C, Supplemental Figure 1A**).

To determine whether aligning to non-TE genomic regions is a special property of siren loci, which produce abundant 24-nt siRNAs in ovules and seed coats, or is a general feature of 24-nt-generating loci from ovule, we compared siRNA abundance and TE coverage in 24-nt windows tiled across all siRNA loci. In ovules, there is a striking negative correlation between read counts and average TE coverage (**Figure 1D**). In contrast, the average TE coverage in leaf is much higher, and only the few exceptional windows with high read count have low TE coverage (**Figure 1E**). Read counts were also plotted against average gene coverage. Ovule read count positively correlated with average gene coverage, with almost 30% of windows overlapping annotated genes (**Figure 1D**), whereas for leaf 24-nt loci only the few exceptional windows with high read count had a similarly high average gene coverage (**Figure 1E**). These results indicate that while siRNA-generating loci frequently overlap TEs, in ovules more siRNA production occurs from non-TE regions of these loci, suggesting that the RdDM pathway in ovule and seed coat may not primarily function to target TEs for silencing. Endosperm siren siRNAs in rice also predominantly map to genic and intergenic regions rather than to TEs (Rodrigues et al., 2013).

### Many siren loci protrude into TEs

Although most siRNA production in ovules arises from non-TE regions, our previous work demonstrated that siren loci are enriched for some classes of TEs (Grover et al., 2020). We therefore analyzed overlap between TEs and siren loci in greater detail. In this study, we defined *B. rapa* siren loci as those loci with >5,000 uniquely-aligning 24-nt reads (157 loci, 1,249-24,379 RPM, **Dataset 1**). While 67% of siren loci overlap an annotated TE, this overlap encompasses only a small fraction of the siren length (8.3%, median) (**Supplemental Table 1, Supplemental Figure 2A**), with TE coverage being generally highest at the edge of the siren while siRNA coverage peaks in the center (**Supplemental Figure 2B**). In contrast, the distribution for the most abundant 24-nt dominant loci in leaves is distinct, with overlapping TEs throughout and peaks in siRNA coverage at the edges (**Supplemental Figure 2C).**This observation suggests that siren loci lie adjacent to, and protrude slightly into, TEs, while the majority of siren siRNAs are produced from regions adjacent to TEs.

When we examined siren loci for overlap with specific families of TEs, two families were found to be significantly enriched - the non-autonomous *Helitron* family *rnd-5_family-1287* (hereafter termed *Persephone)* overlaps 19% of siren loci (z = 55; p = 0; based on 10,000-randomly shuffled windows excluding genes) and the non-autonomous hAT *Bra_hAT1* overlaps 4.5% of sirens (z = 24; p = 0).

The reference sequence used to define the *Persephone* family ends with a short GC-rich hairpin followed 8 bp later by CTAG(T) (**Dataset 2**), as is characteristic of *Helitrons* (Yang and Bennetzen, 2009). It also contains internal repeated sequences, but does not carry exapted gene fragments as some *Helitrons* do. This family has complex structures as a consequence of insertions, truncations, and deletions between pairs of internal repetitive sequences. Despite this variation, alignment of the 27 *Persephone* elements associated with siren loci identified a 74-bp sequence retained in each instance (**Figure 2A, Supplemental Figure 3**), suggesting that this sequence is responsible for siren behavior in ovules. To determine if *Persephone* elements are generally associated with 24-nt siRNA clusters, eight additional *Persephone* family members were examined. Three of these were associated with 24-nt siRNA clusters that did not reach the expression threshold of a siren locus, while five had extremely low siRNA accumulation despite having complete or nearly complete copies of the element (**Supplemental Figure 3**). This observation suggests that the 74-nt sequence might be necessary, but not sufficient, for abundant siRNA production, and that genomic context may also be critical to triggering siRNA hot spots in ovule.

**Figure 2.**
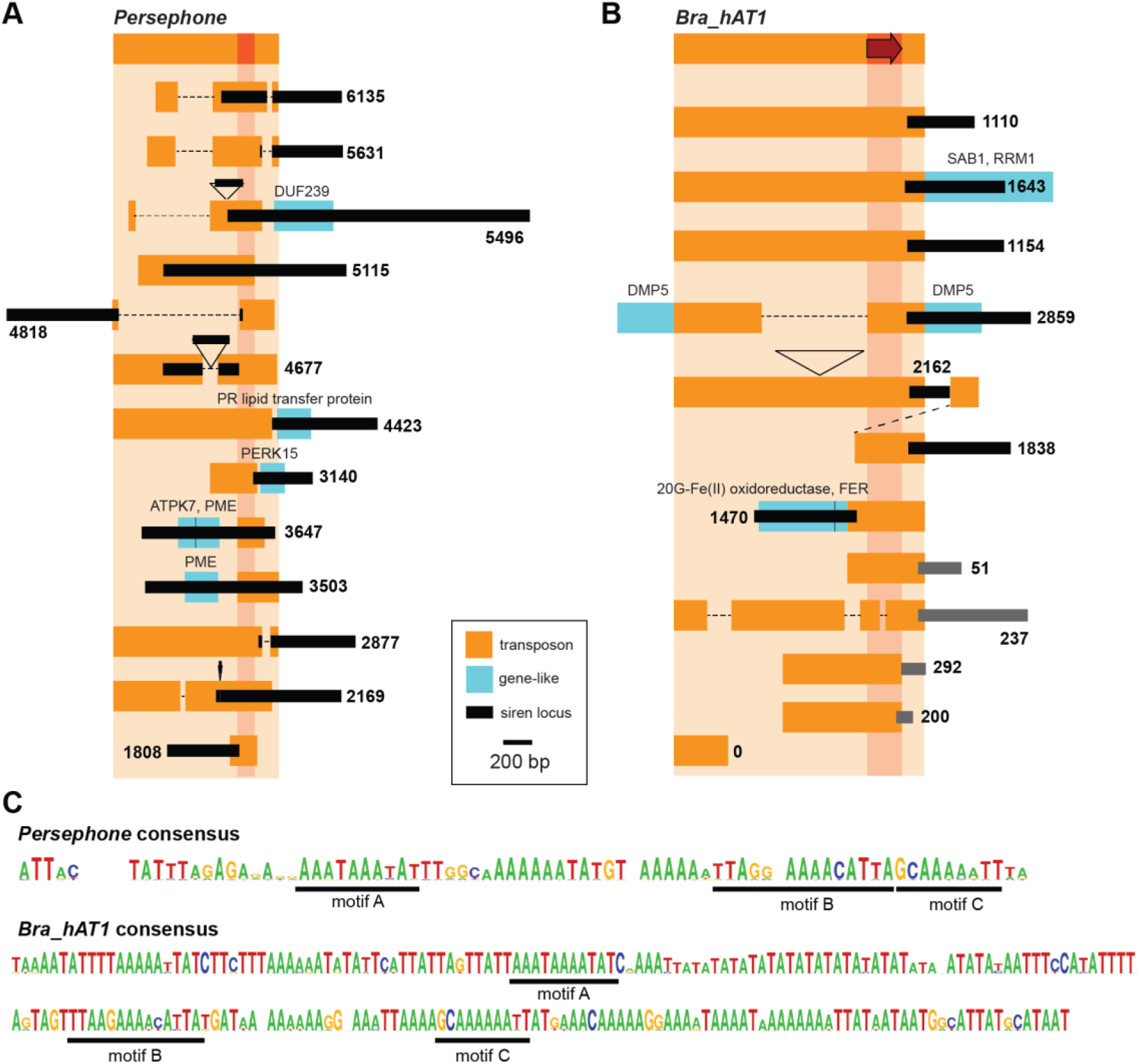
Sirens are associated with non-autonomous *Helitron* and hAT TE families. *Persephone* **(A)**or *Bra_hAT1* elements **(B)**associated with siren loci are shown in orange and are relative to full-length reference sequences (top row). Regions shared by all siren-generating TEs are depicted in dark orange, and an arrow indicates the direction associated with siren locus formation for *Bra_hAT1.* Deletions and insertions are marked by dashed lines and triangles, respectively. A dotted line connects tandem *Bra_hAT1* elements. Siren loci associated with each TE are depicted in black and 24-nt clusters with RPM falling below siren designation are shown in grey; the number of uniquely-aligning 24-nt siRNAs associated with each siren locus is indicated at the side (RPM in ovule). Gene fragments are in blue, labeled with the best match Arabidopsis gene. See Supplemental Figure 3 for additional examples of *Persephone*-associated siren loci. **(C)**Weblogos of the *Persephone* and *Bra_hAT1* regions shared by 24-nt siRNA associated elements. Three motifs shared by these elements are underlined.

We performed similar assessment of the seven *Bra_hAT1*-associated sirens relative to a reference sequence that retains the 8 bp target-site duplication characteristic of this superfamily (**Dataset 2**). Three of the siren-associated *Bra_hAT1* elements are full-length while the others have a variety of internal deletions, insertions, or truncations (**Figure 2B**). In general, siren loci overlap 50-150 nt with one end of the *Bra_hAT1* element, and then extend hundreds of nucleotides in the same direction (see arrow in **Figure 2B**). Only 1 siren locus extends in the opposite direction, but this overlaps the same end of *Bra_hAT1*. To determine whether *Bra_hAT1* is generally associated with 24-nt hot spots in ovule, we used blastn to find the five best additional matches to the *Bra_hAT1* reference sequence. In 4 cases these *Bra_hAT1* elements were associated with 24-nt clusters. In the final case only the end not associated with siren siRNA production is retained, and no 24-nt siRNAs are present (**Figure 2B**). *Bra_hAT1* elements associated with sirens or 24-nt clusters all share an ~200 bp region in common, suggesting that a sequence within this region might be sufficient to trigger 24-nt siRNA expression in flanking sequences.

We looked for possible motifs shared by the 74-bp *Persephone* sequence and the 200-bp *Bra_hAT1* sequence. Runs of adenine are present in both sequences, which is consistent with the presence of A/T rich sequences flanking Pol IV-transcribed regions (Li et al., 2015). Three specific sequence motifs are also present in the same order in each sequence (**Figure 2C**). Motifs B and C are adjacent in the 74-bp *Persephone* sequence and duplicated in one member of this family. The average percentage identity for these 3 motifs is higher in the 30 *Persephone* elements associated with sirens and 24-nt clusters (95% for both groups) than for the 5 *Persephone* elements without an associated 24-nt cluster (84%), suggesting that these motifs might function in siren siRNA production.

### Some ovule sirens contain embedded gene fragments

30% of the highest-expressed 24-nt windows in *B. rapa* ovule loci overlap sequences annotated as genes, prompting us to investigate the relationship between ovule siren loci and gene-like sequences. Forty-four of the 157 siren loci overlap features annotated as genes by at least 40 nt, although comparative analysis indicates that many of these genes are substantially shorter and missing exons, suggesting they are gene fragments (**Supplemental Table 1**). To determine if other siren loci overlap unannotated gene fragments, we used the siren sequence as query in blastn and tblastx searches.

Fifty-two of the 157 siren loci returned Arabidopsis or *B. rapa* genes at an e-value less than 10^-08^(**Supplemental Table 1**), and generally, the peak in siren read count corresponds to the embedded gene fragment (**Supplemental Figure 4**). To determine if siren loci overlap expressed genes, we analyzed ovule mRNAseq data (Grover et al., 2018). Only one siren locus had mRNA expression above 1 FPKM, and most had no evidence of expression (**Supplemental Table 1**). We also defined ovule siren loci in Arabidopsis (Grover et al., 2020) (**Dataset 3**). Twenty-nine of 65 (43%) overlap at least one annotated non-TE gene, including long non-coding RNAs and annotated pseudogenes (**Supplemental Table 2**). Because pseudogenes are often unannotated, we also overlapped Arabidopsis siren loci with a list of 4771 pseudogenes defined by Zou *et al.* (Zou et al., 2009). Twenty siren loci overlapped with pseudogenes from this list (31%), a highly significant number since on average only 3.7 shuffled regions overlap pseudogenes (10,000 randomly shuffled windows; z=8.9; p=0). These results demonstrate that a substantial subset of ovule siren loci in *B. rapa* and Arabidopsis encompass genes, pseudogenes, or gene fragments, and that the gene-like sequences within these loci produce the bulk of siren siRNAs.

### *Ovule siren siRNAs act in* trans *at homologous genes*

Two *B. rapa* siren loci contain gene fragments related to *AT4G03930* (encoding PME42), a pectin methylesterase (PME) gene expressed during early silique development (Louvet et al., 2006) (**Figure 3A**, **Supplemental Figure 4**). Both siren loci also contain truncated *Persephone* elements, and one also carries a fragment related to *AT3G27580,* encoding a protein kinase that phosphorylates an auxin efflux carrier. In both siren loci the majority of 24-nt siRNAs overlap with the gene fragments and not the TE (**Figure 3B**, **Supplemental Figure 4**). These gene fragments have elevated CHH and CHG methylation in ovules relative to leaves, and this methylation requires siRNA production by RDR2 (**Figure 3C-D**). In contrast, methylation in the CG context is high in both ovule and leaves, and is not affected in *rdr2-2* (**Figure 3C-D**). Inspection of DNA methylation over the most closely related full-length PME gene in *B. rapa (A01p015570_BraROA)* revealed high cytosine methylation in all sequence contexts specifically in the region with homology to the siren gene fragments (**Figure 3E-H**). Only methylation in the CG context is present in leaves and *rdr2-2* mutants, and it is present at a reduced level (**Figure 3G**), indicating that methylation at this gene is dependent on 24-nt siRNAs expressed in the ovule. To determine whether siRNAs produced from the two siren loci target methylation at the full-length gene, siren 19-26mer siRNAs were re-aligned to *A01p015570_BraROA* allowing up to two mismatches. 7.96% of reads from one siren locus (608 RPM) and 3.53% of reads from the other siren locus (270 RPM) realigned to *A01p015570_BraROA,* none of them perfectly (**Figure 3F**). In contrast, only 32 siRNAs (5 RPM) originate from *A01p015570_BraROA,* and almost half of these are 22-nt siRNAs. Taken together, these observations suggest that siRNAs produced by the siren-associated gene fragments target methylation at the PME gene in *trans.*

**Figure 3.**
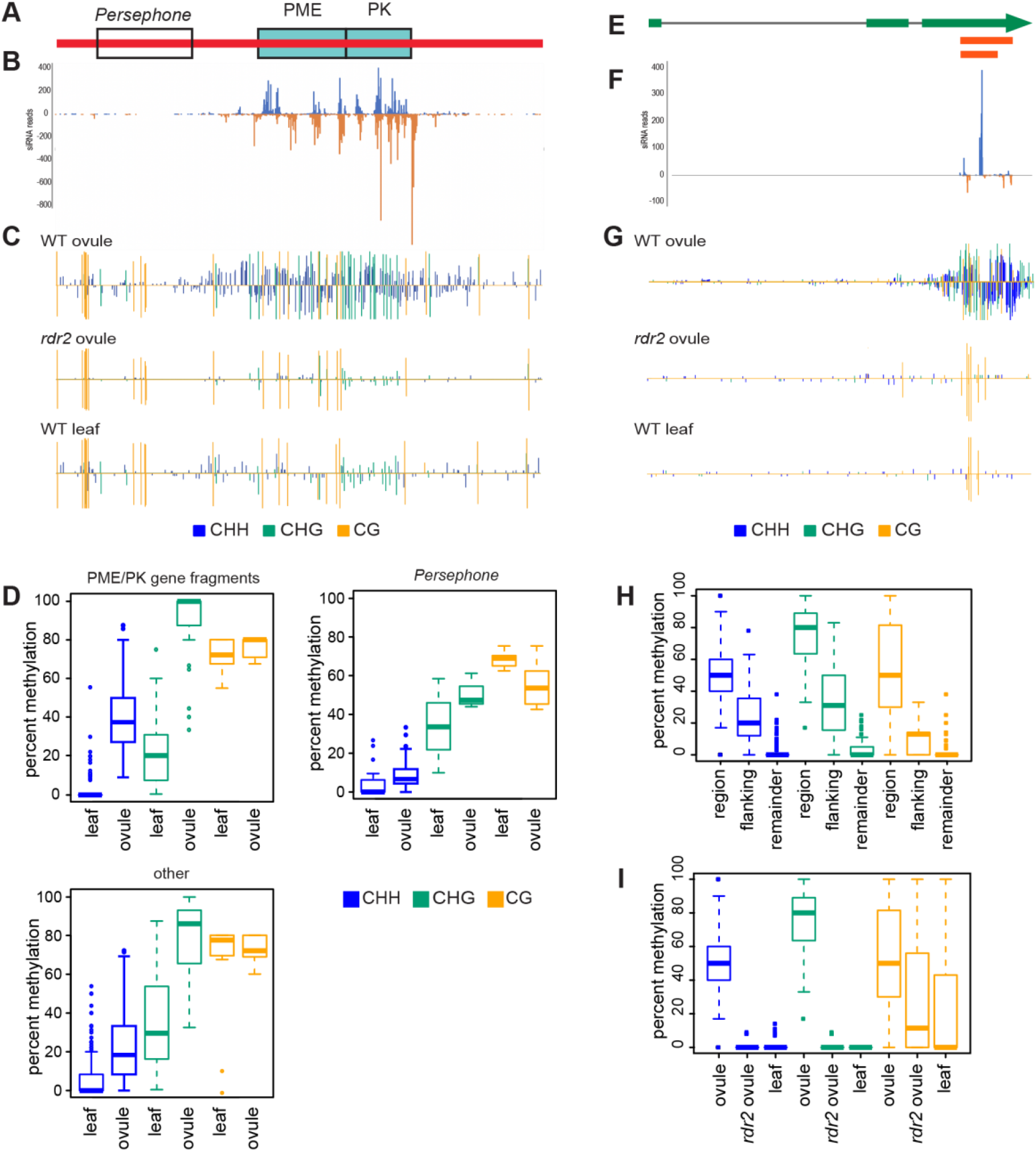
Siren siRNAs *trans*-methylate a protein-coding gene. **(A)**Depiction of the siren (in red) that overlaps a *Persephone* fragment and two gene fragments. **(B)**Distribution of siRNA 5’ ends along the positive strand (blue) and negative strand (orange). **(C)**Methylation along the siren in wild-type ovule, *rdr2-2*ovule, and wt leaf. **(D)**Boxplot of percent methylation of different regions of the siren in leaf and ovule. Only cytosines overlapped by at least 5 reads are included. **(E)**Schematic of the PME gene *A01p015570_BraROA.* HSPs to gene fragments from two different sirens are shown in red. **(F)**Distribution of siRNAs mapping with 1-2 mismatches; there are no perfectly re-aligning siRNAs at this gene. **(G)**Cytosine methylation levels across *A01p015570_BraROA* in ovule, *rdr2-2* ovule, and leaves. **(H)**Boxplots of DNA methylation levels for different regions of *A01p015570_BraROA* (the region of siren homology, the 100 bp region flanking the region of siren homology, and the remainder of the gene). **(I)**Boxplots of DNA methylation for the region of siren homology in ovule, *rdr2-2* ovule, and leaves. Only cytosines overlapped by at least 5 reads were plotted.

A number of other protein-coding genes display elevated non-CG methylation specifically in regions of siren homology identified by BLAST (**Supplemental Figure 5**). This is likely to be *trans*-methylation induced by siren siRNAs, as the RPM of siRNAs originating from the region of the gene (as assessed from uniquely-aligning reads) is low (**Supplemental Table 3**). To identify putative *B. rapa trans-*regulatory targets genome-wide, we re-aligned siren-derived 23-24-nt siRNAs to a TE-masked genome, allowing up to 2 or 3 mismatches between the siRNA and target locus. Windows with at least 1 RPM re-aligning reads and <10% perfectly-matching reads were identified as putative *trans*-methylation sites, resulting in 6,678 and 59,675 windows for 2 and 3 mismatches, respectively (**Supplemental Tables 4-5**). The number of realigning 24-nt siRNAs is strongly correlated with the methylation level, particularly for CHH and CHG methylation (**Figure 4A** and **Supplemental Figure 6A**). When siRNAs with up to 3 mismatches are considered, methylation is lower relative to re-aligning siRNAs, such that only windows with the highest number of re-aligning siRNAs show elevated methylation (**Supplemental Figure 6A-D**). This pattern suggests that siRNAs with 3 mismatches are capable of directing *trans*-methylation, but are less efficient than siRNAs with fewer mismatches. Therefore, our subsequent analysis is limited to windows from the ≤2 mismatch dataset.

**Figure 4.**
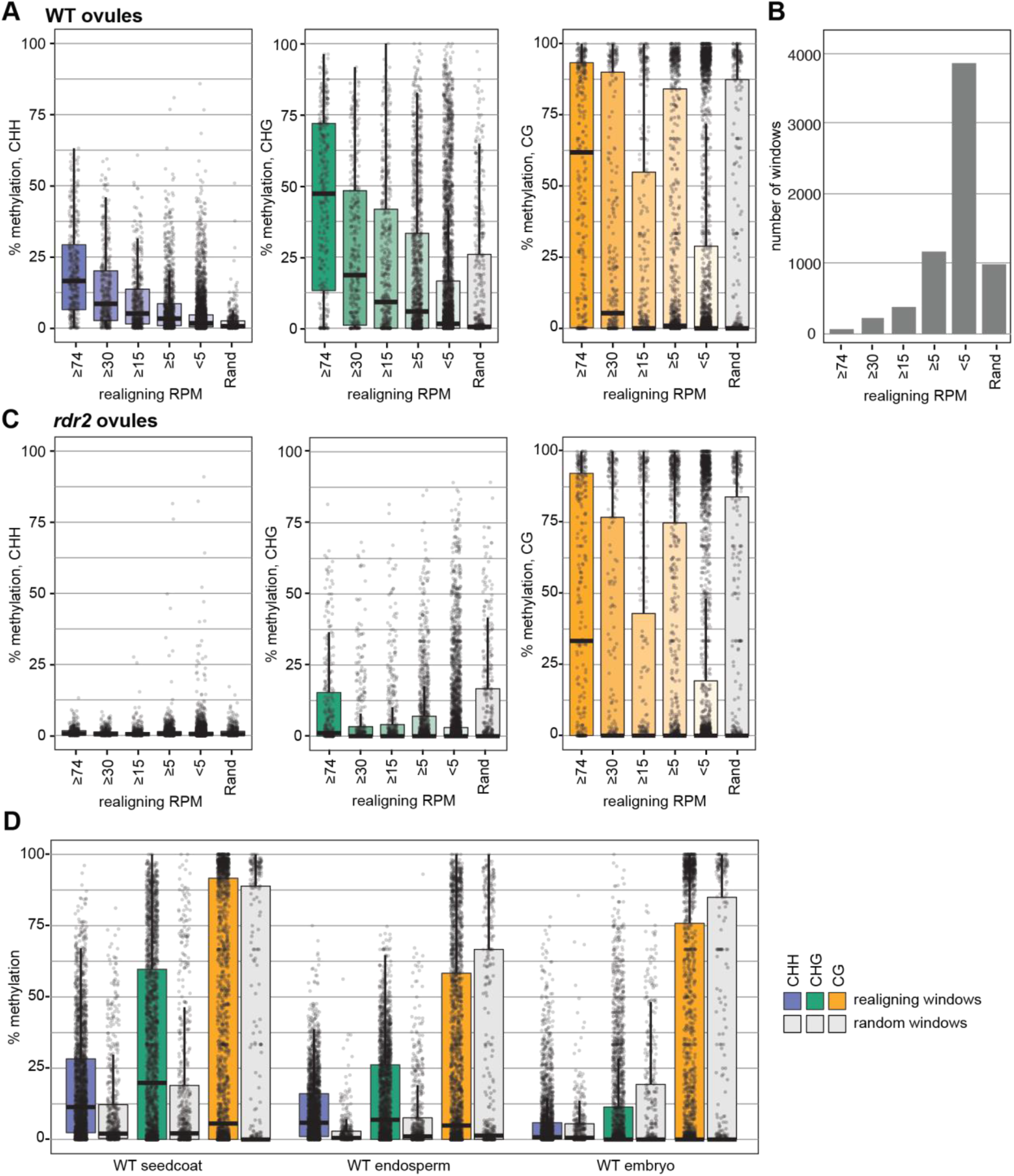
Siren siRNAs *trans*-methylate throughout seed development. **(A)**Box/whisker plots of DNA methylation level at putative *trans-*methylation targets (100-nt genomic windows with ≥1 RPM re-aligning siren siRNAs, ≤2 mismatches) or random genomic windows. **(B)**Bar chart showing the number of windows at each level of re-aligning siRNAs in panels A and C. **(C**) Methylation at putative *trans-*methylation windows in *rdr2* ovules. **(D)**Box/whisker plots for putative *trans-*methylation windows with ≥5 RPM realigning siRNAs in dissected tissues from mid-torpedo stage seeds. For all panels, boxes depict the interquartile range, median values are shown with a black bar, and whiskers encompass datapoints at ≤1.5 times the interquartile range.

To confirm production of siren siRNAs is necessary for *trans*-methylation, we calculated methylation at siRNA re-aligning windows in *rdr2* ovules, which lack 24-nt siRNAs. There is a strong reduction in methylation (**Figure 4C**), indicating that synthesis of 24-nt siRNAs is required for *trans*-methylation of these windows. To eliminate the possibility that the methylation in WT ovules is due to siRNAs produced in *cis,* we filtered our windows to those with zero perfectly re-aligning reads and no production of non-siren reads. Over 90% of windows (6,256) passed this filter, and these exhibit a similar correlation between methylation and siren siRNA realignment (**Supplemental Figure 6E**), indicating that methylation at these sites requires siRNA production at other loci.

After fertilization, siren siRNAs are expressed most highly in the developing seed coat, with moderate accumulation in endosperm and low levels in the embryo (Grover et al., 2020). To investigate whether siren siRNA *trans*-methylation continues after fertilization, we assessed methylation in the three constituent tissues of developing seeds (**Figure 4D** and **Supplemental Figure 7**). Consistent with known accumulation of siren siRNAs, we detect strong *trans-*methylation in developing seed coats and endosperm, with little evidence for siren *trans*-methylation in embryos. These observations suggest that *trans*-methylation occurs throughout seed development in tissues with substantial siren siRNA accumulation.

### Trans-acting siRNAs influence expression of some targeted genes

To investigate the impact of siren siRNA-mediated *trans*-methylation on gene expression, we first assessed expression of the genes with strong homology to siren loci (BLASTn e-value < 10^-08^). Many of these genes are upregulated in *nrpd1* ovules or seeds, suggesting that *trans*-methylation might result in transcriptional silencing (**Supplemental Table 3**). To assess the impact of *trans*-methylation genome-wide, we analyzed the 2,294 siRNA re-aligning windows that overlap an annotated gene by at least 80%. In *nrpd1* ovules, 89 of the 1,270 overlapped genes (7%) are up-regulated and a further 89 genes (7%) are down-regulated (adjusted p-value <0.05, **Supplemental Table 6, Supplemental Figure 8A**). However, many of these genes display only a low level of *trans-*methylation. Target genes with higher CHH and CHG *trans*-methylation are more likely to be upregulated when siren siRNA production is eliminated (**Figure 5AB**). A similar trend toward greater misexpression at higher *trans-*methylation is observed in *rdr2* ovules, but not in *rdr2* leaves (**Supplemental Figure 8B-D**). For 15 candidate upregulated genes we also measured expression in developing seeds via qRT-PCR (**Supplemental Table 7**). Seven target genes were >2-fold upregulated (p < 0.05) in *nrpdl* or *rdr2* 10 days after fertilization (**Figure 5C**). Together, these observations indicate that for some target genes *ŕrans-*methylation is associated with reduced transcript accumulation.

**Figure 5.**
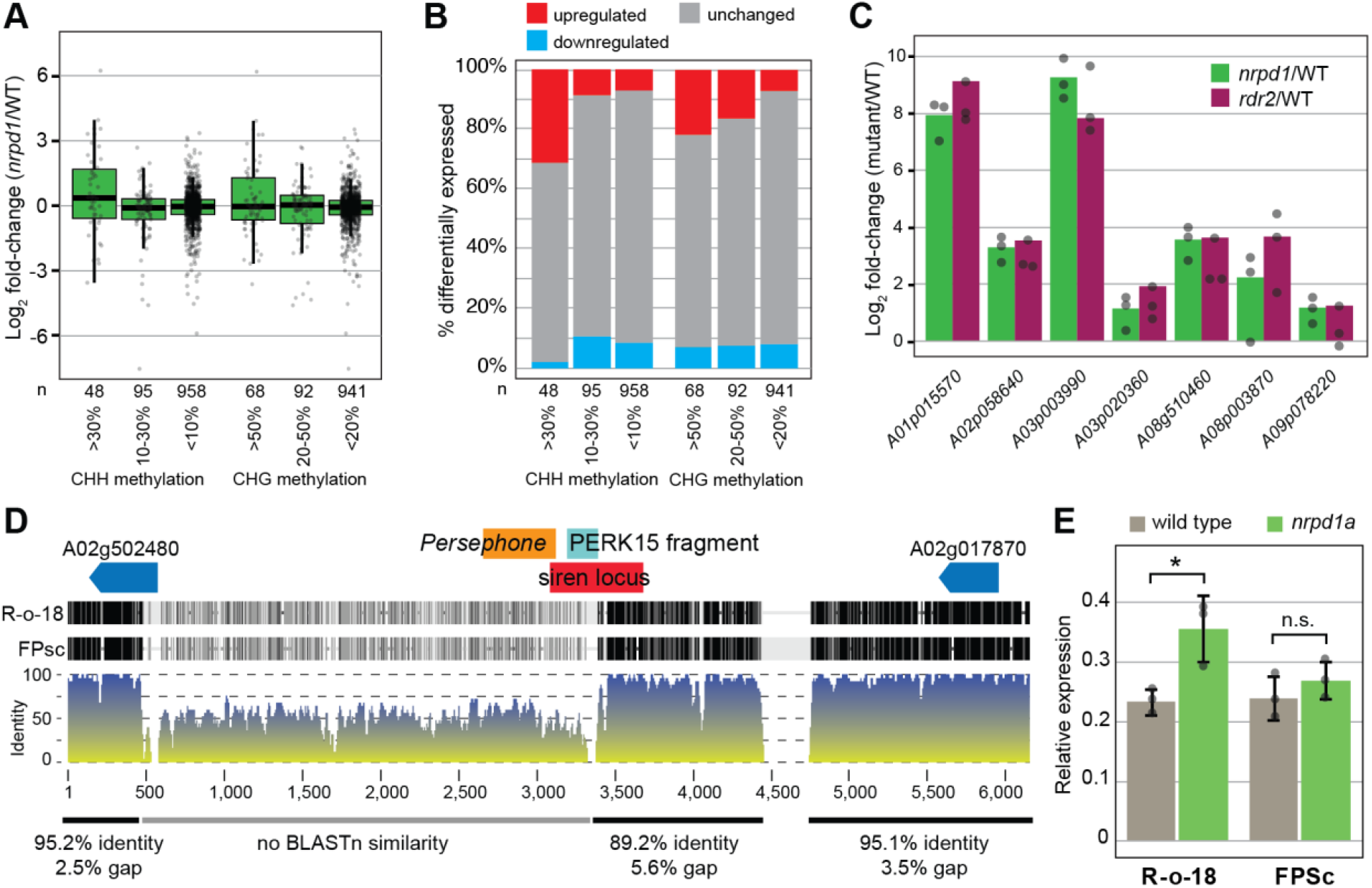
Siren *trans*-methylation impacts gene expression. **(A)**Log_2_ fold-change (*nrpd1*/wild type) of ovule gene expression for the 1270 genes overlapping siren *trans*-methylation sites, stratified by methylation level. Genes with no detectable expression in either genotype were omitted from the analysis. **(B)**Genes from (A) were categorized as upregulated or downregulated when they have an adjusted p-value <0.05. All other genes are considered “unchanged”. **(C)**Fifteen genes that are differentially-expressed in *nrpd1* ovules were tested for differential expression in *rdr2* and *nrpd1* seeds at 10 days post fertilization by qRT-PCR (**Supplemental Table 7**) and the seven genes that are significantly changed are plotted. Bars depict the average log2 fold change versus wild type, error bars are standard deviation of three independent biological replicates. Individual replicates are displayed as dots. **(D)**Alignment of the genomic regions of R-o-18 and FPsc surrounding a polymorphic siren (A02:8131917-8132500). Syntenic genes are shown with blue arrows while components of the siren locus and adjacent TE are labelled as colored boxes. A rolling average of nucleotide % identity is presented below a shaded alignment (black bars indicated shared nucleotides). Identity in distinct regions is summarized below. **(E)**qRT-PCR comparing PERK15 *(BraA08p002810)*expression relative to actin in wild-type and *nrpd1* ovules in the R-o-18 and FPsc backgrounds.

To further test the connection between siren siRNA production and differential expression of homologous genes, we identified a siren locus that is absent in *B. rapa* variety FPsc (**Figure 5D**). SiRNAs from this siren re-align to *PERK15 (BraA08p002810),* which shows *trans*-methylation (**Supplemental Figure 5**) and 1.4-fold upregulation in *nrpdl* RNAseq **(Supplemental Table 6).**We introgressed the *nprd1a* mutation from R-o-18 into FPsc and tested expression of *PERK15* in WT and *nrpd1* ovules of both backgrounds. In R-o-18, which contains the siren locus, *PERK15* is upregulated in *nrpdl* (**Figure 5E**), consistent with our RNAseq analysis. In contrast, there is no difference in *PERK15* expression between WT and *nrpd1* in FPsc where the siren locus is absent. This observation links the production of siRNAs from a siren locus to reduced expression of a homologous gene, demonstrating that siren siRNAs can be *trans-*regulators of protein-coding genes.

### *Fragments from specific gene families are shared by siren loci in* B. rapa *and Arabidopsis*

*B. rapa* and Arabidopsis diverged approximately 14.5 million years ago (mya). This was followed by a whole genome triplication of the *Brassica* genus lineage 10.3 mya and subsequent extensive loss of duplicate genes (Cheng et al., 2013). We previously reported that there is limited synteny between Arabidopsis and *B. rapa* siren loci and no shared sequence between syntenous sirens (Grover et al., 2020), demonstrating the rapid evolution of siren loci. However, we have now found that while syntenic sirens share no sequence in common, they can be related to each other through the retention of non-overlapping gene fragments from an ancestral gene. For a quartet consisting of an Arabidopsis siren locus that is syntenically conserved at all three homeologous positions in *B. rapa* (Grover et al., 2020), the three *B. rapa* sirens contain exons of a DUF239 (neprosin peptidase domain) gene, while the syntenic Arabidopsis siren contains a different exon corresponding to the same gene (**Figure 6AB**). In *Arabidopsis lyrata,* a full-length DUF239-family gene exists in this position, suggesting that this gene was independently converted into a siren locus in *B. rapa* and Arabidopsis. Since the ancestral gene is no longer present in the *B. rapa* genome, and these siRNAs do not align with other DUF239 *B. rapa* genes, it may be that these siren loci have been syntenically conserved to maintain epigenetic structure at this position rather than for *trans-*methylation. However, there is no change in expression of proximal genes when siren siRNA production is eliminated, and therefore it remains unknown why siren character would have convergently evolved at this position.

**Figure 6.**
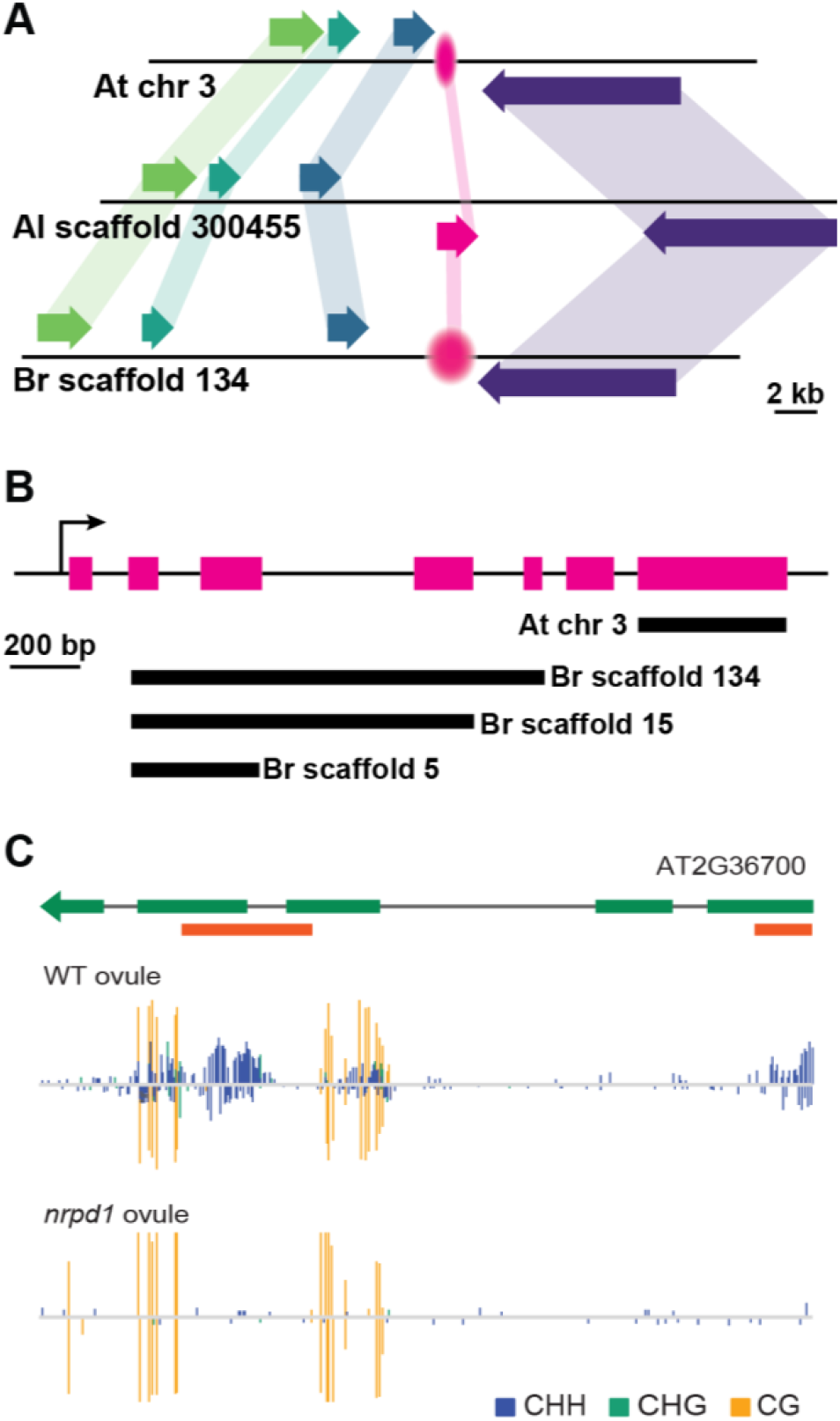
Repeated evolution of siren loci. Siren loci related to DUF239-family genes evolved independently in Arabidopsis and *B. rapa.* **(A)**A syntenic region between *Arabidopsis thaliana* (At), *Arabidopsis lyraŕa* (Al), and *Brassica rapa* (Br). Protein-coding genes are depicted as colored arrows and siren loci as fuzzy ovals. The DUF239 gene is shown in pink. **(B)**Structure of the *A. lyrata*DUF239 gene from (A) with non-overlapping regions of similarity to the *A. thaliana* and three homeologous *B. rapa* sirens shown as black bars. **(C)**A PME gene is also targeted by siren siRNA *trans*-methylation in Arabidopsis. Orange bars beneath the gene model represent regions of homology to siren loci.

Multiple non-syntentic siren loci in Arabidopsis and *B. rapa* also contain fragments of genes from the same gene families, and frequently from the same members of these gene families (**Table 1**, **Supplemental Tables 1-2**). For example, sixteen *B. rapa* and six Arabidopsis siren loci carry fragments from DUF239 domain genes, including three Arabidopsis siren loci and seven *B. rapa* siren loci that are most similar to the same DUF239 gene, *AT5G18460.* In addition to the two siren loci described above that *trans*-methylate PME gene *A01p015570_BraROA,* six additional *B. rapa* siren loci and three Arabidopsis siren loci overlap PME gene fragments. Although the PME family contains 66 members in Arabidopsis, gene fragments found in Arabidopsis and *B. rapa* siren loci are closely related to only 6 members of this family, which themselves fall into two groups of related sequences (Louvet et al., 2006).

**Table 1.** Siren loci in *B. rapa* and Arabidopsis contain fragments homologous to the same gene families.

| Gene family/domain | Number of siren loci with homologous fragments |  | Number of genes most-similar to siren gene fragments |  |
| --- | --- | --- | --- | --- |
|  | *B. rapa* | Arabidopsis | *B. rapa*<sup>a</sup> | Arabidopsis |
| TMK2 receptor-like kinase | 9 | 2 | 2 (1) | 1 |
| Pectin methylesterase | 8 | 3 | 7 (6) | 3 |
| DUF239-containing | 16 | 6 | 6 (3) | 2 |

<sup>a</sup> Number of homologous Arabidopsis genes in parentheses.

To determine whether *trans*-methylation by siren siRNAs is conserved, we assessed ovule methylation at the three Arabidopsis PME genes with similarity to siren gene fragments. Two of these *(AT1G11590* and *AT2G36710)* have methylation in all cytosine contexts across the entire gene and no change in methylation in *nrpd1* ovules. In contrast, *AT2G36700* shows strong *NRPD1-*dependent CHH and CHG methylation specifically in the region homologous to siren loci (**Figure 6C**). RNAseq data demonstrates that *AT2G36700* is upregulated 77-fold in *nrpd1* ovules, 4.6 fold in *clsy3* ovules, but is unchanged in *nrpd1* or *clsy3* leaves (Zhou et al., 2021). *B. rapa* gene *A01p015570.1_BraROA,* a PME targeted by siren *trans*-methylation, is upregulated 75-fold in *nrpd1* ovules and 64-fold in *rdr2* ovules, suggesting that siren-mediated *trans*-methylation represses transcription of PME genes in both Arabidopsis and *B. rapa.* Multiple non-syntenic siren loci targeting the same genes or gene families in Arabidopsis and *B. rapa* suggests convergent evolution of siren siRNA-mediated *trans*-methylation.Alternatively, conserved siren loci might be rapidly repositioned in the genome.

### Tissue-specific expression of siren siRNAs requires CLSY3 function

To investigate factors responsible for expression of siren loci, we overlapped Arabidopsis ovule siren loci with 24-nt siRNA clusters from unopened flower buds of various siRNA mutants (Zhou et al., 2018). As expected, all 65 Arabidopsis siren loci overlap *nrpd1-*dependent loci. Additionally, 58 of the siren loci overlap *CLSY3*-dependent loci and the remaining 7 siren loci overlap 24-nt siRNA loci that were affected by the *clsy3 clsy4* double mutant (**Supplemental Table 2**). Loci dependent on *CLSY1, CLSY2*, or *SHH1* had no overlap with siren siRNA loci. Zhou *et al* have also established that siRNA production in ovules is dependent on *CLSY3* and *CLSY4*, which are highly expressed in ovules (Zhou et al., 2021).

We therefore examined transcript accumulation of the CLASSY gene family in *B. rapa.* Following the Brassica whole genome triplication, one gene has been retained in *B. rapa* for *CLSY1 (A08p019180.1_BraROA), CLSY2 (A10p025980.1_BraROA),* and *CLSY3 (A10p004670.1_BraROA),*and two genes for *CLSY4 (A07p011690.1_BraROA, A03p046540.1_BraROA).* As in Arabidopsis, *CLSY3* is the predominant family member expressed in ovule, with both *CLSY1* and *CLSY2* expressed at very low levels (**Figure 7A**). *CLSY3* is also the predominant family member expressed in 10 dpf seed and endosperm. Expression of *CLSY3* is much higher than expression of *CLSY4* in both ovule and endosperm, which is consistent with the stronger requirement for *CLSY3* for siRNA production at siren loci in Arabidopsis (Zhou et al., 2021). In tissues lacking substantial siren siRNA accumulation, such as leaf and embryo, *CLSY3* expression is low and *CLSY1* is the predominant family member expressed.

**Figure 7.**
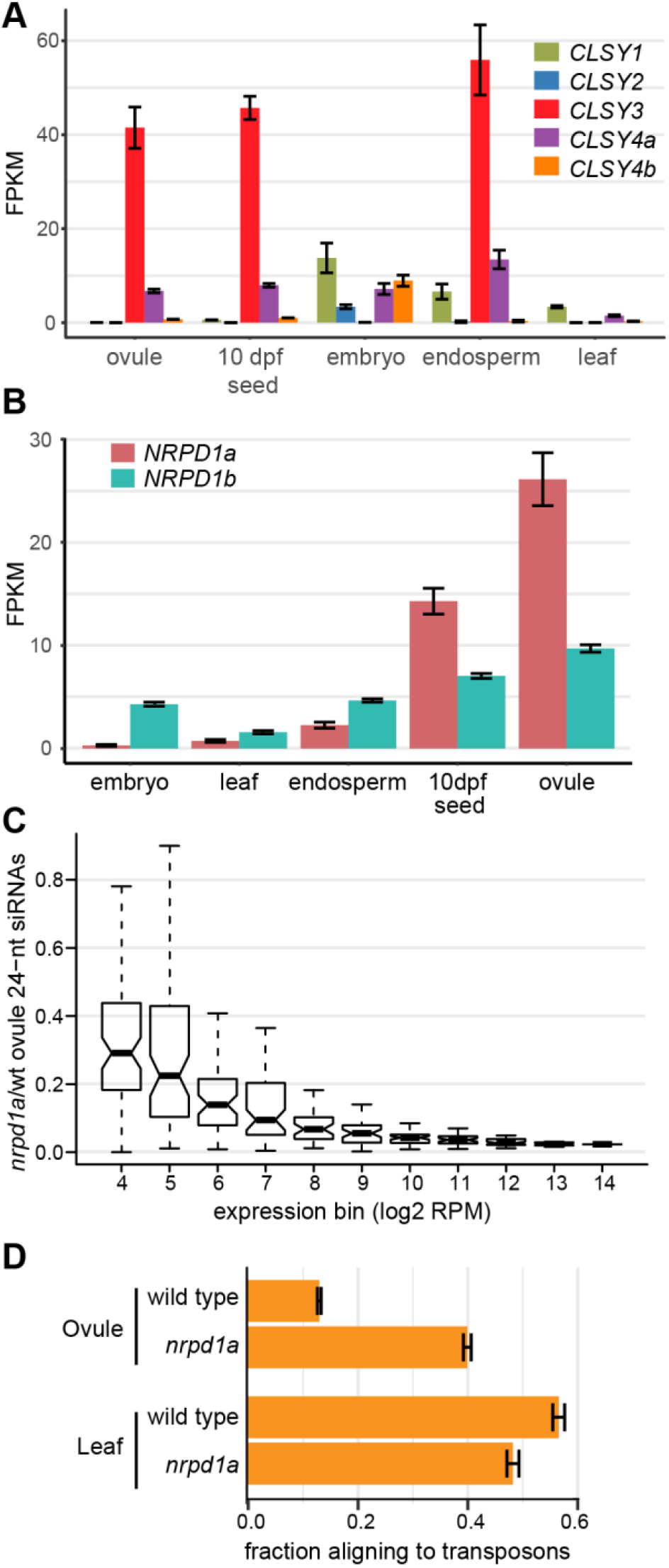
***CLSY3, CLSY4,* and *NRPD1a* are associated with ovule-specific expression in *B. rapa.*** Transcript accumulation of CLSY genes **(A)**and NRPD1 orthologs **(B)**in *B. rapa*. Error bars show the standard error of the mean from three RNAseq replicates. **(C)**Ovule loci producing abundant 24-nt siRNAs are more strongly impacted by the *nrpd1a* mutation. Outliers are not shown. **(D)**The fraction of 24-nt siRNAs aligning to TEs increases in the ovule *nrpd1a* mutant, but is unchanged in leaves. Error bars depict the standard error of the mean of three sRNA-seq

In Arabidopsis, an 18-bp sequence is enriched at sites of CLSY3 binding (Zhou et al., 2021). We compared this sequence with the conserved motifs from *Persephone* and *Bra_hAT1* elements. When compared to the adjacent motifs B and C in the *Persephone* sequence, the reverse complement of the CLSY3-binding sequence is identical at 14 of 19 positions (74%, **Supplemental Figure 9**). It is therefore possible that *Persephone* and *Bra_hAT1* elements carry sequences associated with CLSY3 binding, allowing them to induce siRNA production specifically and abundantly in ovules.

### *A paralog of NRPD1 has specialized for siren siRNA production in* B. rapa

We previously showed that 24-nt siRNAs from ovule are almost completely lost from a *B. rapa rdr2* mutant (Grover et al., 2018). However, some siRNAs remain in the *B. rapa nrpd1-2* mutant, which has a missense mutation in the highly conserved Metal A binding site (**Supplemental Figure 10A-B)** (Haag et al., 2009; Huang et al., 2013). While earlier *B. rapa* genome assemblies included only one *NRPD1* gene, an improved genome assembly includes a second *NRPD1* gene, providing a potential source for Pol IV activity in the *nrpd1-2* mutant. The two *NRPD1* paralogs are differentially expressed (**Figure 7B**), with ovule expressing relatively more of the copy corresponding to *nrpd1-2 (BraA.NRPD1a, A09p015000.1_BraROA),* while leaves and the filial tissues (embryo and endosperm) predominantly express the second copy *(BraA.NRPD1b, A08g503970.1_BraROA).*We therefore investigated whether *NRPD1a* might be specifically responsible for production of the abundant non-TE siRNAs in ovules. **Figure 7C** shows that there is a positive correlation between expression level of an siRNA locus in wild type and loss of siRNA accumulation in *nrpd1-2.* In contrast, no such correlation exists for *rdr2* or *nrpe1* (**Supplemental Figure 10C**). This correlation suggests that *NRPD1a* is required for expression of the most abundant siRNA loci in ovules. We also compared the fraction of siRNAs arising from TE or non-TE sequences and discovered that *nprd1-2* ovules contain a TE-rich population of siRNAs that is similar to the siRNA population in leaves (**Figure 7D**), indicating that these TE-enriched loci are not influenced by *NRPD1a.* Together these observations suggest that *NRPD1a* specifically impacts expression of siren siRNAs in *B. rapa* ovules, while *NRPD1b* might be either redundantly or solely responsible for expression of canonical RdDM loci.

## DISCUSSION

The canonical RdDM pathway is well-described in plants as a pathway for control of TEs (Matzke and Mosher, 2014; Stroud et al., 2013). Here we show that the population of 24-nt siRNAs generated via this pathway differs significantly between organs. In leaves most 24-nt siRNAs originate from TEs, in accordance with the cooperative role they play in maintaining TE silencing. In ovule a much smaller percentage of 24-nt siRNAs map to TEs largely due to the abundant population of 24-nt siRNAs generated from siren loci. Production of siRNAs from siren loci is associated with expression of the putative chromatin remodeler CLSY3 and one of two copies of NRPD1. Siren siRNAs are frequently associated with gene fragments and can induce CHH and CHG methylation in *trans* at related genes despite mismatches between the siRNA and target region. siRNAs with ≤2 mismatches are associated with greater methylation than siRNAs with 3 mismatches, although the latter are associated with *trans*-methylation when present in sufficient numbers. It is likely that the position of mismatches across the siRNA is an important factor for its function (Chipman and Pasquinelli, 2019). The recent observation of widespread Pol V transcription (Tsuzuki et al., 2020) provides a plausible mechanism for siRNA-mediated recruitment of methyltransferases to target genes in *trans,* although Pol II might also create a scaffold for these siRNAs to function (Zheng et al., 2009).

In maize, some TEs contain gene fragments from “donor” protein-coding genes, resulting in siRNAs that perfectly match both the TE and the donor gene. Such genes have higher methylation relative to non-donor genes (Muyle et al., 2021); however, since the origin of such siRNAs is ambiguous, it is unclear whether this methylation is due to *trans*-methylation or due to production of siRNAs at the protein-coding gene. Either way, the implication is that when TEs contain sequences captured from protein-coding genes, the host risks “collateral” silencing of donor genes. Here, we demonstrate methylation of protein-coding genes by *trans*-action of 24-nt siRNAs, including siRNAs that do not perfectly match the target gene (**Supplemental Figure 6E**). The abundant production of siRNAs from gene fragments, rather than from the neighboring TE sequences (**Figure 3, Supplemental Figure 4**), suggests that activity at the protein-coding gene is more than a collateral effect during reproductive development.

Siren-like loci that produce highly abundant 24-nt siRNAs were recently described in Arabidopsis tapetum, somatic cells surrounding the male germline (Long et al., 2021). It is not reported whether these loci carry gene fragments, however siRNAs produced from these sites function in *trans* to trigger methylation and silencing. Like the ovule siren siRNAs described here, siRNAs from abundant tapetal loci also require CLSY3 (Zhou et al., 2021; Long et al., 2021), raising the possibility that similar siren-induced *trans*-methylation functions in both maternal and paternal somatic tissue surrounding the germline. On the paternal side, siRNAs are produced in tapetum and trigger methylation in the male meiocyte. We have previously proposed that siren siRNAs accumulating in the endosperm might be produced from maternal integuments (Grover et al., 2020; Chakraborty et al., 2021). Here, we show that siren targets are methylated in the endosperm, however it remains to be tested where these siRNAs originate. Importantly, the ability to function despite mismatches between the siRNA and the target locus indicates that maternally-derived siren siRNAs could function on both maternally- and paternally-derived alleles in endosperm. Imbalance between the dosage of maternal siRNAs and paternally-derived targets could contribute to maternal- or paternal-excess seed phenotypes (Lu et al., 2012). *Trans*-methylation at closely related sequences also implies that a single siren locus could impact diverging homeologous sequences following whole genome duplication.

Tapetal siren-like siRNAs silence transcription of a *Gypsy* retrotransposon through *trans*-methylation of its LTR (Long et al., 2021). We also detected changes in expression for some of the *trans*-methylated genes in *nrpd1a* ovules and young seeds (**Figure 5AB, Supplemental Table 3**). Gene body methylation in the CG context occurs most often in moderately transcribed genes, but its effect, if any, on transcription is controversial (Bewick et al., 2016; Zilberman et al., 2007). Genes with heavy methylation tend to be more upregulated in a *met1* mutant, suggesting that methylation within gene bodies could impede transcript elongation (Zilberman et al., 2007). However, unlike the *trans*-methylated genes in this study, gene body methylation is absent from the first 2 kb and last 1 kb of genes (Zilberman et al., 2007). The *trans*-methylated genes in this study also differ in being highly methylated in CHH and CHG contexts, and while CG methylation of genes has been shown to correlate with expression, CHG methylation anticorrelates with expression (Schmitz et al., 2013). In *Arabidopsis lyrata,* genes gaining gene body CHG methylation in endosperm are associated with reduced gene expression, and the magnitude of increase in gene body CHG methylation in maternal alleles positively correlates with increased expression bias in favor of the paternal allele (Klosinska et al., 2016). Although changes in expression of siren target genes are most likely a consequence of DNA methylation, we cannot yet eliminate the possibility that siren siRNAs might induce post-transcriptional gene silencing in parallel with DNA methylation.

We particularly noted that three gene families are associated with multiple sirens in both *B. rapa* and Arabidopsis: PMEs, TMK receptor-like kinases, and DUF239 genes (**Table 1**). In each case the same phylogenetic group within these families is associated with siren loci, further suggesting functional significance of siren association. A reduction of only 18% in PME activity in a pollen-specific isoform is sufficient to affect pollen tubes, resulting in reduced fertilization (Jiang et al., 2005). Thus, siren-generated siRNAs could play a critical role in regulating the expression of PME genes, including the regulation of multiple closely-related family members within this large family. Siren loci may also play some additional unknown function as shown by the existence of a set of siren loci that are syntenically conserved in Arabidopsis and all three *B. rapa* homeologous positions that are related to each other through an ancestral DUF239 gene, but have no sequence homology to each other and no longer have strong sequence homology to any DUF239 genes existing in their own genome.

Whether their function is primarily *trans-*methylation or something else, hints of the importance of siren siRNAs come from the phenotype of the *B. rapa nrpd1-2* mutant, which has a severe seed abortion phenotype (Grover et al., 2018). *nrpd1-2* ovules lack siren siRNAs but produce most canonical 24-nt siRNAs (**Supplemental Figure 10AB**), indicating that seed production might rely on siren siRNAs in *B. rapa.* Interestingly, the *nrpd1-2* phenotype is determined by the maternal sporophytic genotype (Grover et al., 2018), supporting a model whereby maternally-produced siRNAs cause *trans*-methylation in the endosperm to enable its proper development (Kirkbride et al., 2019).

Some siren loci appear to derive from the pseudogenization of a once functional gene, based on the presence of a likely functional copy at a tandem position or at a syntenic position in close relatives. Although we observed a correlation between non-functional gene fragments and siren loci, we cannot determine which characteristic is causal. RdDM might enable initial pseudogenization or might be recruited by non-coding pseudogenic transcripts. Other siren gene fragments appear to be insertions relative to Arabidopsis, perhaps formed by the addition of filler DNA during double-strand break repair, as has been proposed for gene fragments captured by *Helitrons* (Kapitonov and Jurka, 2007).

Although a few siren loci are centered on a TE, most siren loci overlap a TE at the edge of the locus and not in the region where the majority of 24-nt siRNAs are produced (**Supplemental Figure 2**). Two specific families of TEs, the *Helitron Persephone* and the non-autonomous *Bra_hAT1* element, are associated with almost a quarter of siren loci. For both TEs, additional family members were also associated with 24-nt siRNA clusters, suggesting that these elements may include motifs capable of inducing the production of siRNAs in adjacent sequences in the ovule. Alternatively, the siren environment might be favored for insertion by these elements. However, we believe this less likely because for both families, tandem copies of the TE resulted in the formation of independent sirens. In addition, for both TEs, internal deletions amongst family members reduced the relevant sequence to a very small region (74 bp for *Persephone* and 200 bp for *Bra_hAT1).* A *Bra_hAT1* element retaining the wrong end of the element was the only family member not associated with 24-nt clusters. The region of these elements that is associated with siren siRNA production includes sequences similar to a CLSY3 binding motif (**Supplemental Figure 9**), suggesting a novel mechanism for TEs to produce epigenetic variability.

TEs have been exapted for a number of functions (reviewed in (Lisch, 2013)), including providing regulatory elements, placing genes under epigenetic control, and serving as the source of new protein-coding genes. Here we show that certain TEs may be able to induce large amounts of siRNAs to be made from adjacent sequences in an organ-specific fashion. SiRNAs arising from these regions cause *trans*-methylation at, and in some cases impact expression of, closely-related genes. Such regulation could account for the sporophytic requirement of RdDM for seed development in *B. rapa.*

## MATERIALS AND METHODS

### Biological material

*Brassica rapa* subsp. trilocularis, R-o-18 genotype, was used for all experiments unless otherwise noted. Unfertilized ovules were collected less than 24 h prior to anthesis. *B. rapa* RdDM mutants were obtained from a TILLING population, then backcrossed six times to the parental line, as previously described (Grover et al., 2018). RdDM mutant lines *braA.nrpd1.a-2, braA.rdr2.a-2,* and *braA.nrpe1.a-1* are referred to as *nrpd1a, rdr2,* and *nrpe1,* respectively. All gene names correspond to the publicly-released v2.3 genome (NCBI GCA_017639395.1).

FPsc *nrpd1* mutants were generated by crossing to *braA.nrpd1.a-2* and backcrossing five times to wild-type FPsc. Genotyping primers (5’-CGGAGAACGAGAGTTTCAAGCAG-3’ and 5’-CGCATAAACAACTCGAGCGTCT-3’) were used to confirm that the polymorphic siren (A02:8131917-8132500) is missing in FPsc *nrpd1* individuals.

### Analysis of TEs

*B. rapa* TEs were annotated as previously described (Grover et al., 2018), with the addition of RepBase (Bao et al., 2015) *Brassica* and Arabidopsis repetitive elements updated through vol 20, issue 3 and ten *B. rapa* TRIM elements (Gao et al., 2016). The TEs used in the annotation also included 1519 TEs from a RepeatModeler-generated dataset (Cheng et al., 2016); 1005 of these have been classified to family, 658 of which we have manually-annotated to generate full-length exemplars (**Dataset 4**). Detailed methods for TE annotation are in the **Supplemental Methods**. Masked *B. rapa* and Arabidopsis genomes were generated by running RepeatMasker (v. 4-0.5, A.F.A. Smit, R. Hubley& P. Green, www.repeatmasker.org) under- nolow option to exclude masking of simple repeats. The combined library of *Brassica* and Arabidopsis repetitive DNA elements resulting in masking of 41.5% of the *B. rapa* R-o-18 v2.3 genome and 20.25% of the Arabidopsis v10 genome.

*Bra_hAT1* and *Heliŕron rnd-5_family-1287 (Persephone)* elements were aligned to the full-length exemplar sequences (**Dataset 2**) using Muscle (https://www.ebi.ac.uk/Tools/msa/muscle/) (Edgar, 2004), with the alignment manually corrected based on blastn alignments between each individual family member and a full-length exemplar.

### Small RNA alignment

*B. rapa* 19-26 bp small RNA reads were retrieved from the NCBI SRA (BioProject PRJNA588293). Wild-type samples arose from hand-dissected samples of leaves, unfertilized ovules immediately before anthesis, or embryos, endosperm, and seed coat dissected from mid-torpedo stage seeds. Seed coats were further bisected and separated into “chalazal” (near the funiculus) and “upper” (distal from the funiculus). Small RNA reads were aligned to *B. rapa* genotype R-o-18 v2.3 using Bowtie (Langmead et al., 2009) under conditions permitting only perfectly-matched unique alignments (-v 0, -m 1) or under conditions in which perfectly-matched reads multiply-aligning up to 49 times are positioned by ShortStack (Johnson et al., 2016) based on local densities of uniquely-aligning reads (--mismatches 0, --mmap u). RPM calculations were based on the number of alignable 19-26 nt siRNAs in a library and mutant libraries were normalized based on 21-nt small RNA read number. Arabidopsis small RNA reads (NCBI SRA SRR5646727-SRR5646729) were processed in parallel.

To re-align siren siRNAs, 23-24 nt Shortstack-aligned reads at siren loci were captured from the bamfile using the BEDTools intersect command (Quinlan and Hall, 2010), converted into a fastq file using the samtools fastq command (http://www.htslib.org/) (Li et al., 2009), and realigned to a genome masked for TEs and sirens using bowtie but allowing up to 2 or 3 mismatches (-v 2, -v 3). The number of realigning reads and perfectly aligning reads was parsed from the data and quantified in 100-nt windows tiled every 50-nt. The number of small RNA reads originating from genes was determined by intersecting perfectly-matched and uniquely-aligning ovule small RNA reads with *trans*-methylated gene targets, or the region of these genes with blastn matches to sirens (HSP region), using BEDTools intersect.

### siRNA locus analysis

24-nt siRNA ovule or leaf reads were size-selected and clustered using BEDTools v2.25.0 merge and unix commands. Overlapping 24-nt siRNA clusters with at least 10 reads were merged when separated by no more than 100 bp. For the comparison of 24-nt siRNA abundance with TE and gene coverage, sliding windows with a step size of 12-nt were generated across ovule and leaf 24-nt clusters generated using Shortstack-aligned reads. The fraction coverage of each window by TEs or genes was determined using the bedtools coverage command. The number of 24 nt siRNAs in each cluster was determined using the bedtools intersect command, requiring at least half of the siRNA to be present in each window (-c -F 0.5).

*B. rapa* 24-nt clusters with at least 5000 uniquely-aligning reads (1245 RPM) were defined as siren loci, and an equivalent RPM was used to define the 65 Arabidopsis ovule siren loci (**Datasets 1, 3**).These siren loci are similar but not identical to previously reported ovule “core” siren loci (Grover et al., 2020). For a positional profile of 24-nt siRNA 5’ ends mapping across sirens, a bedfile of 24-nt reads aligned to each siren using ShortStack was generated using the BEDTools intersect command; for each strand the number of 5’ ends at each position was counted using a custom perl script.

Overlap between siren loci and genomic features (TEs, genes, pseudogenes, CLSY-dependent loci) was determined using the BEDTools intersect command. To determine the statistical significance of overlaps with pseudogenes and TEs, 10,000 sets of non-overlapping genomic intervals of matching size were generated using BEDTools shuffle (-noOverlapping) and intersected with the pseudogene or TE feature. Genes were excluded from matching genomic intervals for the TE shuffle.

TE coverage across siren loci was averaged over 100 bp windows starting at each end and preceding in 50 bp steps; at least five siren loci were included in each averaged window. TE coverage in each window was calculated using the BEDTools coverage command. The average fraction of siren 24-nt siRNAs in each 100 bp window was determined using the BEDTools intersect command.

Syntenic orthologs between Arabidopsis and *B. rapa* were identified as previously described (Grover et al., 2020). The best-matching Arabidopsis and *B. rapa* genes (e-value < 10^-08^) were identified with blastn and tblastx (performed using Blastall) with siren sequences or genes overlapped by siren sequences as queries. Genomic regions carrying siren loci were compared to genomic regions carrying best hit Arabidopsis and *B. rapa* genes with GEvo (https://genomevolution.org/CoGe/GEvo.pl) (Lyons et al., 2008) using genomes that had been annotated for siren loci and TEs.

### Chromosomal distribution analysis

24-nt uniquely-aligning siRNA clusters with at least 10 overlapping reads were mapped to R-o-18 v2.3 pseudochromosomes in 100,000 bp windows. A TE-masked genome was generated using RepeatMasker. Pericentromeric and large gene-poor heterochromatic regions were inferred using *Gypsy* retrotransposon coverage. The BEDTools coverage command was used to determine the fraction of each window covered by *Gypsy* retrotransposons. Leaf read counts were normalized to ovule read counts based on the number of alignable 19-26 nt sRNAs in the libraries.

### DNA *trans*-methylation analysis

*B. rapa* methylation data was retrieved from the NCBI SRA (BioProjects PRJNA588293 and PRJNA657007) and aligned to the *B. rapa* v2.3 genome with a WGBS Snakemake workflow utilizing Trim Galore, bwa-meth, and MethylDackel (Grover, 2019). Bisulfite conversion rates, as determined by Lambda DNA, were at least 99%. Arabidopsis ovule methylation (NCBI GEO SRR13404120-SRR13404122, SRR13404131) was visualized with CoGe LoadExp+’s Methylation Analysis Pipeline using Bismark (Grover et al., 2017).

### mRNA-seq data analysis

RNAseq libraries from *rdr2* ovules were prepared with Amaryllis Nucleics RNAseq library kit before sequencing on an Illumina NextSeq. The resulting libraries were deposited at the NCBI SRA (BioProject number PRJNA808098). RNAseq data from wild-type and *nrpd1* ovules were obtained from SRA accession SRP132223 (Grover et al., 2018). Trimmed RNAseq reads were aligned to the R-o-18 v2.3 genome using STAR v2.5.4b (Dobin et al., 2013). Reads overlapping annotated genes were counted using htseq-count (Anders et al., 2015). Replicate consistency was checked by principle component analysis on rlog-transformed counts generated by DESeq2 (Love et al., 2014). Differentially expressed genes (v2.3 annotation) and FDR-corrected p-values were determined using DESeq2.

Differential expression of Arabidopsis PME gene AT2G36700 in *clsy* mutant ovules was retrieved from Zhou et al., 2021.

### qRT-PCR

Three independent replicates of *B. rapa* ovules or 10 dpf seeds were collected for extraction of total nucleic acid (tNA) (White and Kaper, 1989). Replicate samples came from plants grown together in the same greenhouse. 2 μg tNA were DNaseI digested with DNA-free kit DNase Treatment and Removal Reagents (Ambion) following the manufacturers’“rigorous” DNase treatment protocol. After removal of contaminating DNA, 20 μL (1.8 μg) was incubated with 1 μL 50μM random hexamers and 1 μL 10 mM dNTP mix at 65°C for 5 mins, followed by cDNA synthesis with SuperScript IV Reverse transcriptase (Invitrogen) according to the manufacturers’ protocol with the following reaction: 5 μL 5x SSIV buffer, 1 μL 100 mM DTT, 1 μL RNaseOUT RNase Inhibitor (40 U/μL), and 1 μL SuperScript IV Reverse transcriptase.

To quantify gene expression, qRT-PCR was carried out with 1 μL of a 1:1 diluted cDNA, 0.625 μL of each 10 mM gene specific primer (**Supplementary Table 8**), 12.5 μL 2x SensiMix SYBR& Fluorescein master mix (Bioline) and 10.25 μL nuclease-free water. Twenty-four putative *trans-*methylation target genes were selected for analysis; 23 were expressed at a detectable level. The melting curve from each primer set was checked to ensure the specificity of the qRT-PCR reaction and the expression level of each transcript was normalized to *ACTIN2.* Three independent biological replicates were compared to determine fold-change and p-values.

### Accession numbers

Sequence data from this article can be found in NCBI SRA under accession numbers SRR5886891-SRR5886893 (*B. rapa* siRNAs), SRR10415409-SRR10415420 (*B. rapa* WGBS), SRR6675211-SRR6675222, PRJNA808098 (*B. rapa* ovule and 10 dpf RNAseq), SRR5646727-SRR5646729 (Arabidopsis flower bud siRNAs), SRR185818 (Arabidopsis ovule siRNAs), and SRR13404120-SRR13404122, SRR13404131 (Arabidopsis ovule WGBS).

## ACKNOWLEDGEMENTS

We are grateful to Dr. Eric Lyons and the CoGe team for development and maintenance of CoGe. We thank Dr. Damon Lisch for critical reading of an earlier draft and for suggesting that we look for transmethylation. The authors are grateful for support from the National Science Foundation (IOS-1546825 to RAM and MF) and the USDA National Institute of Food and Agriculture (AFRI 2021-67013-33797 to RAM and AFRI 2014-67013-21661, subaward C0471A-B to MF).

## AUTHOR CONTRIBUTIONS

DB, HTC, MF, and RAM designed the research; all authors performed research; DB, HTC, MF, and RAM analyzed data; DB, HTC, MF, and RAM wrote the paper. All authors read and approved the manuscript.

