## Supplemental Materials for "Ovule siRNAs methylate protein-coding genes in *trans*"

**Supplemental Materials for Burgess D, *et al.* 2022**

**Contents:**

Supplemental Methods

Supplemental Tables 7-8

Supplemental Figures 1-10

**Additional files:**

Supplemental Table 1. *B. rapa* sirens

Supplemental Table 2. Arabidopsis sirens

Supplemental Table 3. Curated *B. rapa* target genes

Supplemental Table 4. Re-aligning windows, ≤2 mismatches

Supplemental Table 5. Re-aligning windows, ≤3 mismatches

Supplemental Table 6. Genes overlapping re-aligning windows

Dataset 1: *B. rapa* ovule siren sequences

Dataset 2: Exemplar sequences for *Persephone* and *Bra_hAT1*

Dataset 3: Arabidopsis ovule siren sequences

Dataset 4: Exemplar sequences for *B. rapa* R-o-18 transposon annotation

### Supplemental Methods

#### Transposon annotation in the *B. rapa* R-o-18 genome

To produce full-length exemplars a set of best matching sequences was retrieved and aligned using Muscle (https://www.ebi.ac.uk/Tools/msa/muscle/). These sequences were obtained through a combination of retrieving the five best RepeatMasker-generated matches (v. 4-0.5) based on the Smith-Waterman score or the top ten blastn matches using CoGe BLAST (https://genomevolution.org/CoGe/CoGeBlast.pl). Up to 1 kb of sequence was added at both ends to the RepeatMasker-generated matches using the Bedtools slop command. The top 10 CoGe blast matches were examined on CoGe GEvo panels (https://genomevolution.org/coge/GEvo.pl), and the largest high scoring segment pair (HSP) for each sequence was retrieved and extended by 20-50 nt. From the multiple sequence alignment full-length sequences were obtained and classified to superfamily based on characteristic features (Wicker, 2012) and on matches to reference repeats in Repbase (https://www.girinst.org/censor/). LTR retrotransposons were subdivided into LTR and internal region based on the presence of LTR direct repeats (5'-TG…3'CA) flanked by a 5 bp target site duplication (TSD) and the presence of a polypurine tract at the end of the internal region. LTR direct repeats and DNA transposon terminal inverted repeats (TIRs) were detected by BLASTing sequences against themselves. Non-autonomous TRIM and LARD retrotransposons were classified based on the size of their non-coding internal region (<600 bp and >2000 bp, respectively). LINE and SINE elements were defined based on a 3' poly(A) sequence and a flanking 7-21 bp TSD. DNA transposons were classified based on the TSD and characteristic TIR sequences. The ends of Helitrons were defined based on 5'(A)TC and 3'CTRR(T), and the presence of a small G/C-rich hairpin sequence near the 3' end. Captured host sequences, identified as blastn hits to Arabidopsis CDS sequences with an e-value of 1E-05 or less, were masked with N's to avoid the erroneous identification of genes as TEs by RepeatMasker. RepeatModeler sequences that were not manually annotated were sorted to superfamily based on blastn hits with an e-value of 1E-05 or less to classified TE sequences.

**Wicker, T.** (2012). So many repeats and so little time: How to classify transposable elements. In Plant Transposable Elements, Topics in current genetics. (Springer Berlin Heidelberg: Berlin, Heidelberg), pp. 1–15.

**Supplemental Table 7. qRT-PCR results with p-values from 10 dpf seeds**

| Gene name | Fold change (*nrpd1*/WT) | p-value | Fold change (*rdr2*/WT) | p-value |
|---|---|---|---|---|
| A01p003160.1_BraROA | 0.39 | 6.80E-02 | 0.59 | 1.27E-01 |
| A01p015570.1_BraROA | 247.71 | **8.86E-03** | 344.64 | **1.02E-02** |
| A02p011510.1_BraROA | 1.09 | 9.89E-01 | 2.55 | 3.78E-01 |
| A02p057100.1_BraROA | 1.42 | 3.37E-01 | 2.78 | 6.03E-01 |
| A02p058640.1_BraROA | 9.86 | **1.33E-04** | 8.09 | **7.49E-03** |
| A03p003990.1_BraROA | 622.00 | **1.84E-02** | 404.34 | **4.68E-02** |
| A03p004630.1_BraROA | 1.27 | 4.15E-01 | 0.97 | 8.41E-01 |
| A03p020360.1_BraROA | 2.18 | **4.62E-02** | 2.61 | **2.85E-02** |
| A03p033380.1_BraROA | 2.90 | 3.61E-01 | 0.91 | 5.38E-01 |
| A05g501190.1_BraROA | 0.68 | 9.20E-02 | 0.65 | 2.45E-01 |
| A06p049610.1_BraROA | 1.26 | 4.09E-01 | 0.44 | **8.17E-03** |
| A08g510460.1_BraROA | 11.95 | **4.90E-03** | 7.13 | 5.98E-02 |
| A08p002810.1_BraROA | 1.10 | 7.06E-01 | 1.07 | 8.14E-01 |
| A08p003870.1_BraROA | 4.66 | 6.89E-02 | 12.68 | **2.89E-03** |
| A09p078220.1_BraROA | 2.23 | **2.00E-02** | 1.48 | 3.12E-01 |

[1] p-values in **bold** are significant at 0.05 following FDR correction

### Supplemental Table 8. qRT-PCR primers

| Gene name | F primer (5' to 3') | R primer (5' to 3') |
|---|---|---|
| A03p043420.1_BraROA (ACT2) | CTTGCACCAAGCAGCATGAA | CCGATCCAGACACTGTACTTCCTT |
| A01p003160.1_BraROA | CGACAAACACTAGCGTCGGACA | GTCGCCGTAACTAAATCCGCAA |
| A01p015570.1_BraROA | GAAAGCCTAATCCAGGACAGAGC | TGACAGGAGAAAGATCCGCACT |
| A02p011510.1_BraROA | CATAGTGGCCGAACTCTTGGAC | AAAGGGAAGCCCTGATGGTC |
| A02p057100.1_BraROA | GCAAGACCTTGTTGTGATGCTC | GGAATCCGACAGAGTTCAGGC |
| A02p058640.1_BraROA | CACCACCAGAGGTGATAGAGC | GTTACTGGAGCCGTTCTGTCAC |
| A03p003990.1_BraROA | CGCTTCAAATGTGCTTCATGTGAC | GCTCCACATGAACATTGGAAGACTC |
| A03p004630.1_BraROA | CAACGTCGCGAATCCGACA | CGAGCTTCTGGTCGTCATCG |
| A03p020360.1_BraROA | CTTCTTGGAGTGCAAGGCTGATG | GAAGCCTTGAGGCTCGCTATG |
| A03p033380.1_BraROA | CGGTGAGTGTAACTATAGATACCCAGT | ACGCGTTCTTGCAGTCGTC |
| A05g501190.1_BraROA | GCATGTCGATGAAAGTTGTATCCAG | GCATTGAGAGACAGAACGGTGAC |
| A06p049610.1_BraROA | CACAGCGCCTCTCTACACAG | TCGTTGTTAGCTCTGTGAGGCT |
| A08p002810.1_BraROA | CTAATCCACCGGACCCTGT | GAGGGGTTGAAGATGGTTCGATG |
| A08p003870.1_BraROA | CCTTCATCGGCGTACTTGTGG | GCTGTTGAAAAACTTGAGCCCAGT |
| A08g510460.1_BraROA | GCTCTATGCCCACATTGAGCGA | TGTTCTGTTGCCCCTGGAGT |
| A09p078220.1_BraROA | GCGTTGAGCTTGGAGGAGCA | CAAACAAATCAGCAACGGGACCA |

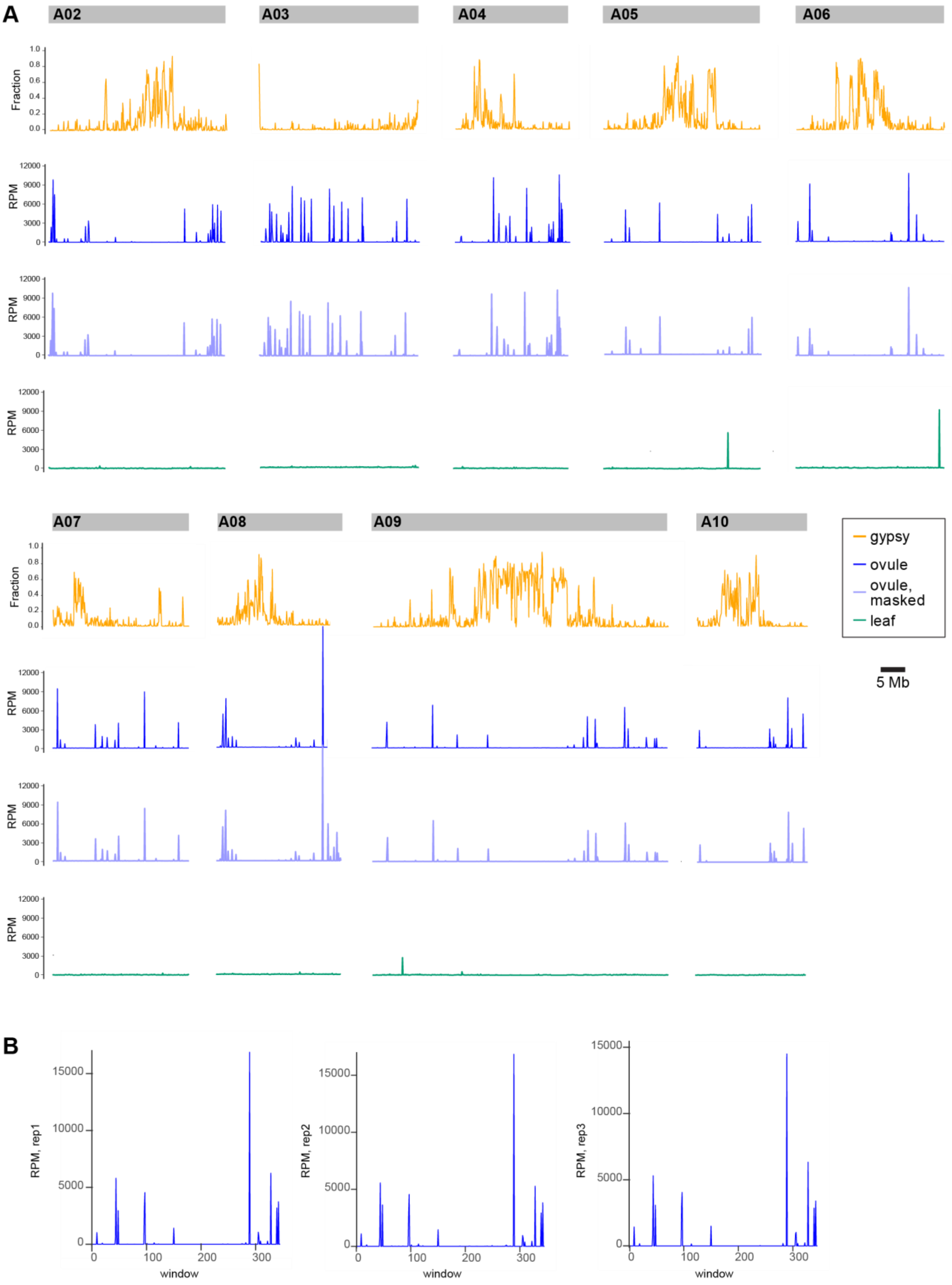
A
A02
A03
A04
A05
A06
A07
A08
A09
A10
Fraction
RPM
gypsy
ovule
ovule, masked
leaf
5 Mb
B
RPM, rep1
RPM, rep2
RPM, rep3
window

**Supplemental Figure 1. *B. rapa* ovule 24-nt siRNAs predominantly map to non-repetitive hotspots on all chromosomes (supporting Figure 1C).**
(**A**) Distribution of pooled, uniquely-aligning ovule and leaf siRNA on *B. rapa* R-o-18 v2.3 chromosomes 2-9 (chromosome 1 shown in Figure 1C). RPM in 100 kb windows is plotted. Discrete, highly-expressed peaks in leaf data correspond to the intersection of sense/antisense transcripts or to transcribed regions with secondary structure. SiRNA production from these regions does not require RDR2 and is therefore not canonical RdDM. (**B**) Replicates of uniquely-aligning ovule siRNA abundance over chromosome 1 demonstrate the consistency of biological replicates.

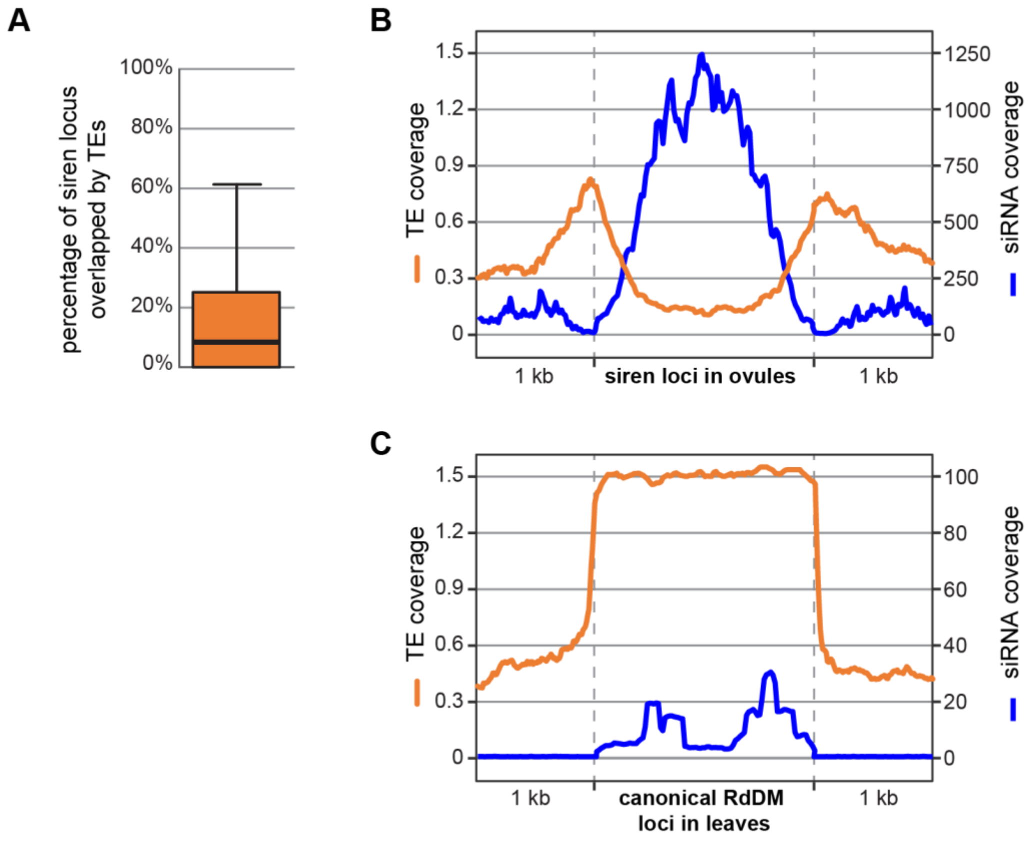


**Supplemental Figure 2. Siren loci overlap TEs primarily at their edges (supporting Figure 1D-E).** **(A)** Distribution of TE coverage over *B. rapa* siren loci. **(B)** Metaplots of TE and siRNA coverage across siren loci **(B)** or the 200 most abundantly-expressed 24-nt dominant loci in leaves (≤100 nt in length) **(C)**.

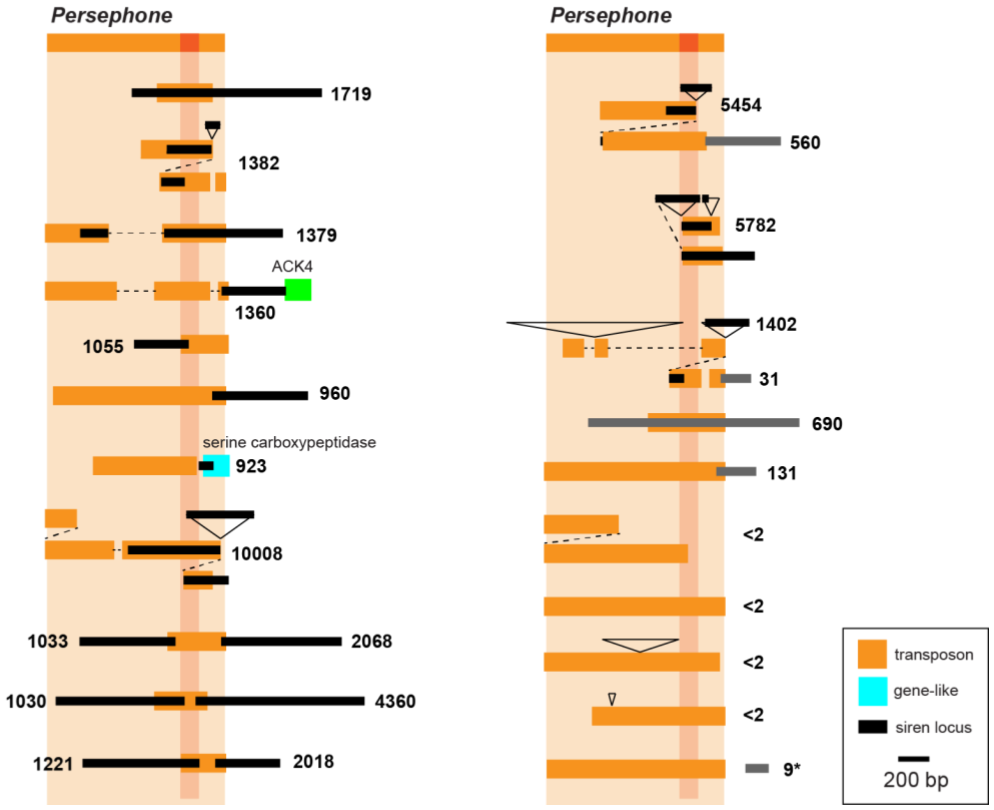


**Supplemental Figure 3. Additional examples of *Persephone* elements (supporting Figure 2).** Siren sequences are shown in black and 24-nt clusters with RPM falling below siren designation are shown in grey. Deletions are marked by dashed lines, insertions by triangles designating the length of the insertion. A dotted line connects tandem *Persephone* elements. The number of uniquely-aligning 24-nt siRNAs in each 24-nt cluster is indicated in RPM (reads per million alignable 19-26mer siRNAs in the pooled ovule libraries). Siren regions containing gene fragments, shown in blue, are labeled with the best hit *A. thaliana* gene. Shown in green is the 3'-UTR and part of the coding region of a gene related to Arabidopsis peroxisomal acyl-CoA oxidase *ACX4* (AT3G51840). Not all gene fragments or overlapping genes are shown. The final *Persephone* element in this figure, delineated with an asterisk, has a low number of 24-nt siRNAs in ovule, but a substantially increased number in seed coat.

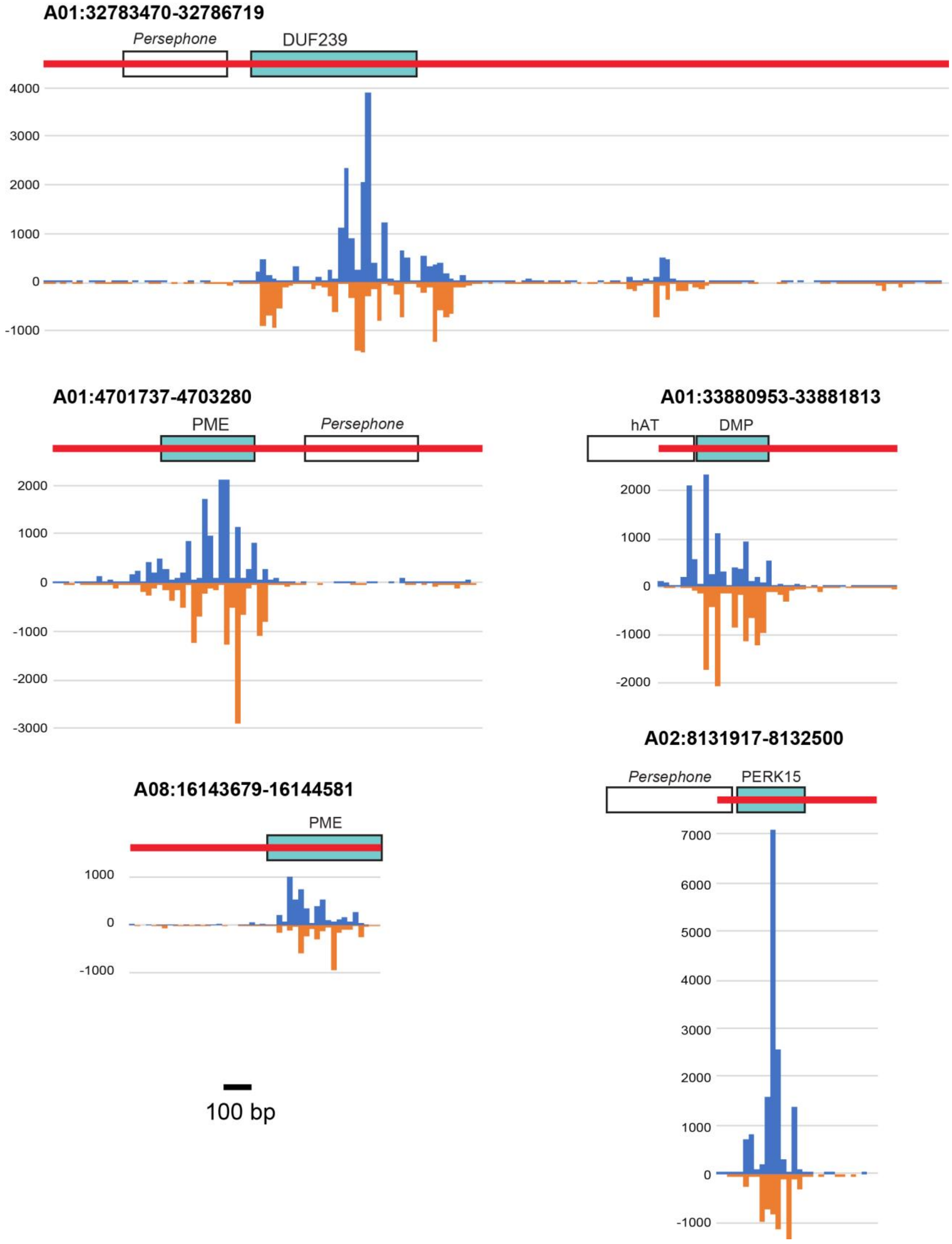
A01:32783470-32786719
Persephone
DUF239
A01:4701737-4703280
PME
Persephone
A01:33880953-33881813
hAT
DMP
A02:8131917-8132500
Persephone
PERK15
A08:16143679-16144581
PME
100 bp

**Supplemental Figure 4. siRNA accumulation peaks in siren gene fragments (supporting Figures 2, 3A).**

Generally, siRNA accumulation is most abundant from gene fragments (blue boxes) within siren loci (red bars). Transposable elements (white boxes) are not the primary source of siRNAs. Total accumulation of siRNA 5’ ends in 20 bp windows tiled across the siren locus are shown below (blue, sense; orange, antisense). The second siren locus containing a gene fragment similar to PME42 (AT4G03930) is A01:4701737-4703280.

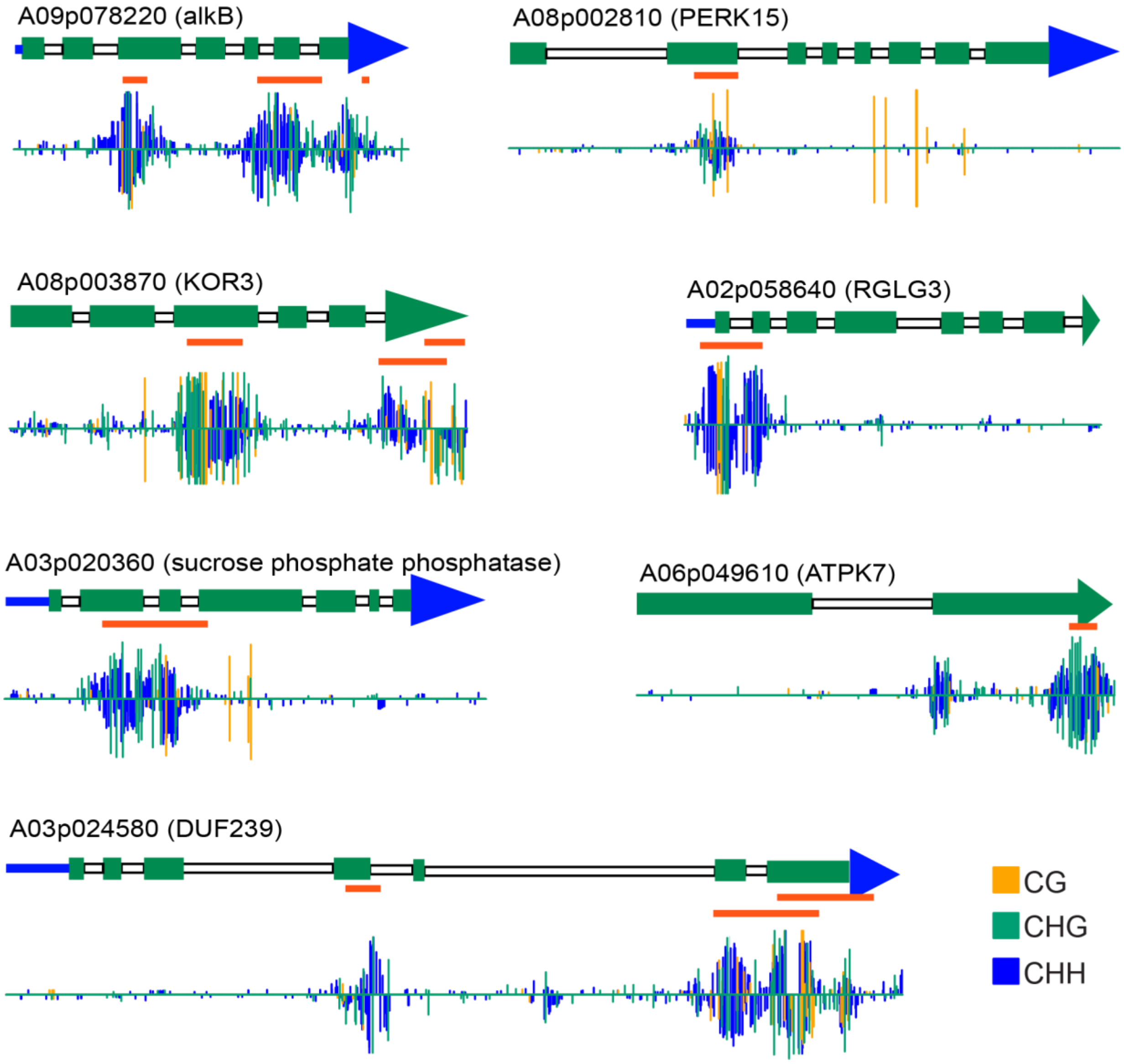


**Supplemental Figure 5. Protein coding genes are specifically methylated at regions of homology to siren gene fragments (supporting Figure 3G).**

Cytosine methylation in ovule for several protein-coding genes with homology to siren gene fragments. The name of each gene is listed above the gene model, with the Arabidopsis homolog listed in parentheses. Coding exons are shown in green, UTRs are shown in blue and introns in white. Regions with similarity to siren gene fragments are depicted with an orange bar beneath the gene model. Below each gene model is a representation of cytosine methylation across the gene, demonstrating that abundant methylation coincides with the region of similarity to the siren gene fragment.

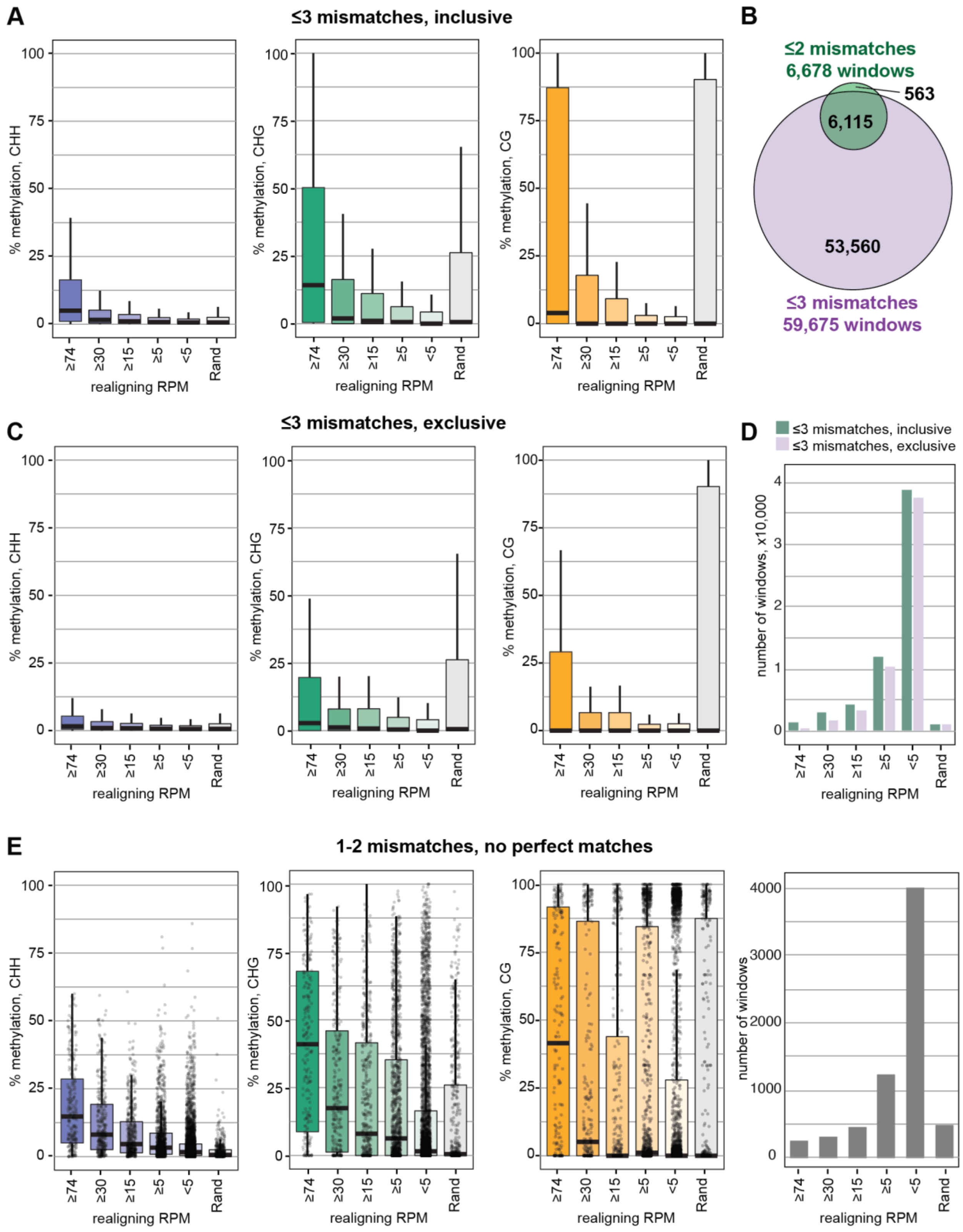


**Supplemental Figure 6. Re-aligning siren siRNAs are associated with increased methylation (supporting Figure 4A).**

**(A)** Methylation at 65,790 genomic windows with >1 RPM when ≤3 mismatches are allowed (purple circle in **(B)**. **(B)** Representative Venn diagram of genomic windows passing the 1 RPM threshold when 2 or 3 mismatches are allowed. 563 windows are found only in the ≤2 mismatch set because some siRNAs are removed from the analysis when they realign to >50 sites when 3 mismatches are allowed. **(C)** 53,560 windows identified only when 3 mismatches are allowed, but not identified when only 2 mismatches are allowed (≤3 mismatches, exclusive), have very little signal of *trans*-methylation. **(D)** Histogram of the number of windows in each re-aligning RPM category in panels A, C. **(E)** 3,323 windows with >1 RPM realigning reads (1-2 mismatches), none of which perfectly match the target window and therefore cannot be produced in *cis*. In all plots, boxes represent interquartile range and whiskers encompass datapoints at ≤1.5 times the interquartile range. Median is marked by a black line.

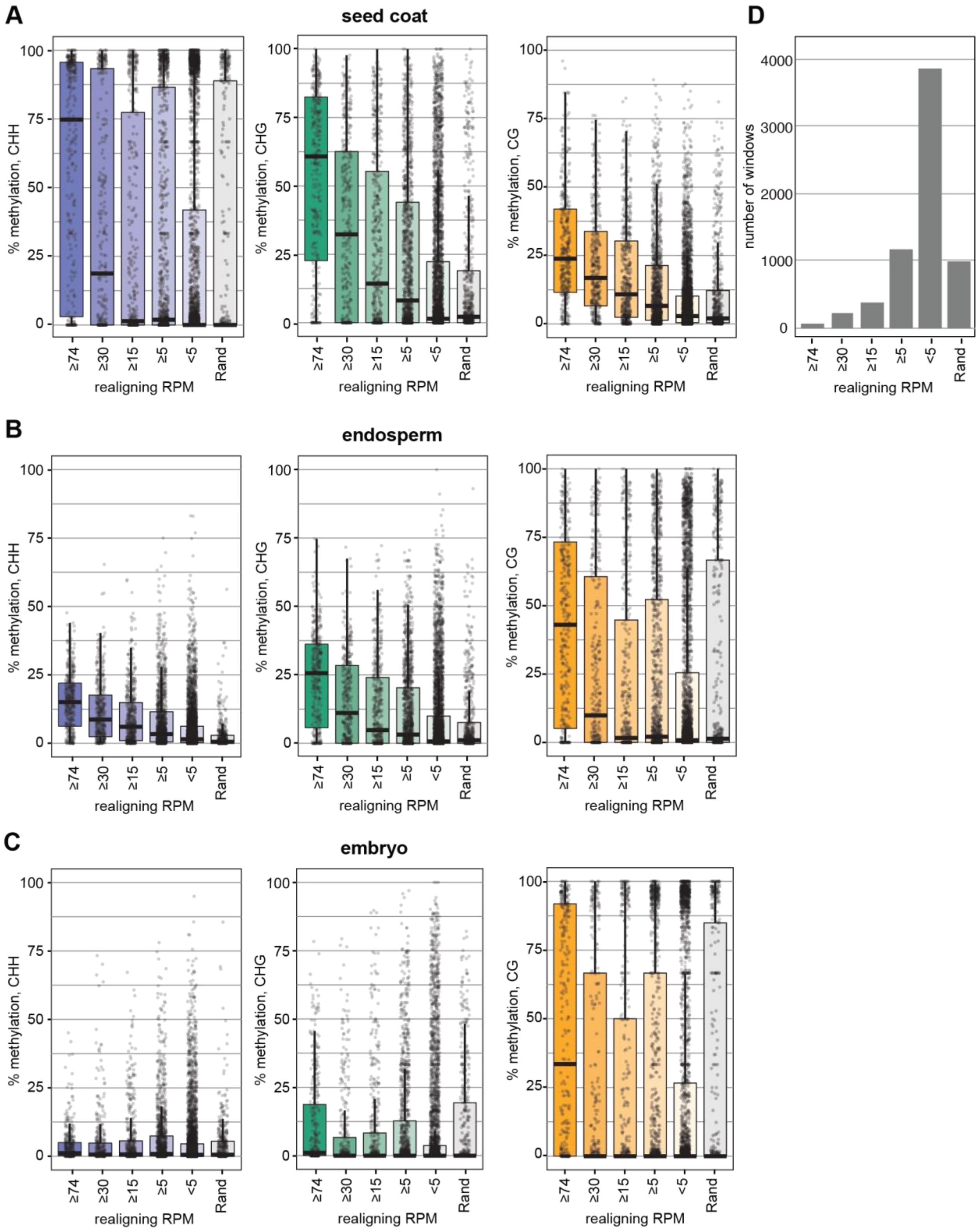


**Supplemental Figure 7. *Trans*-methylation continues post-fertilization in seed coat and endosperm (supporting Figure 4D).**

Methylation at the 6,678 windows from Figure 4A, assessed in the seed coat **(A)**, endosperm **(B)**, and embryo **(C)** broken down by number of realigning reads. Boxes depict the interquartile range, median values are shown with a black bar, and whiskers encompass datapoints at ≤1.5 times the interquartile range. (**D**) Histogram of the number of windows in each re-aligning RPM category in panels A-C.

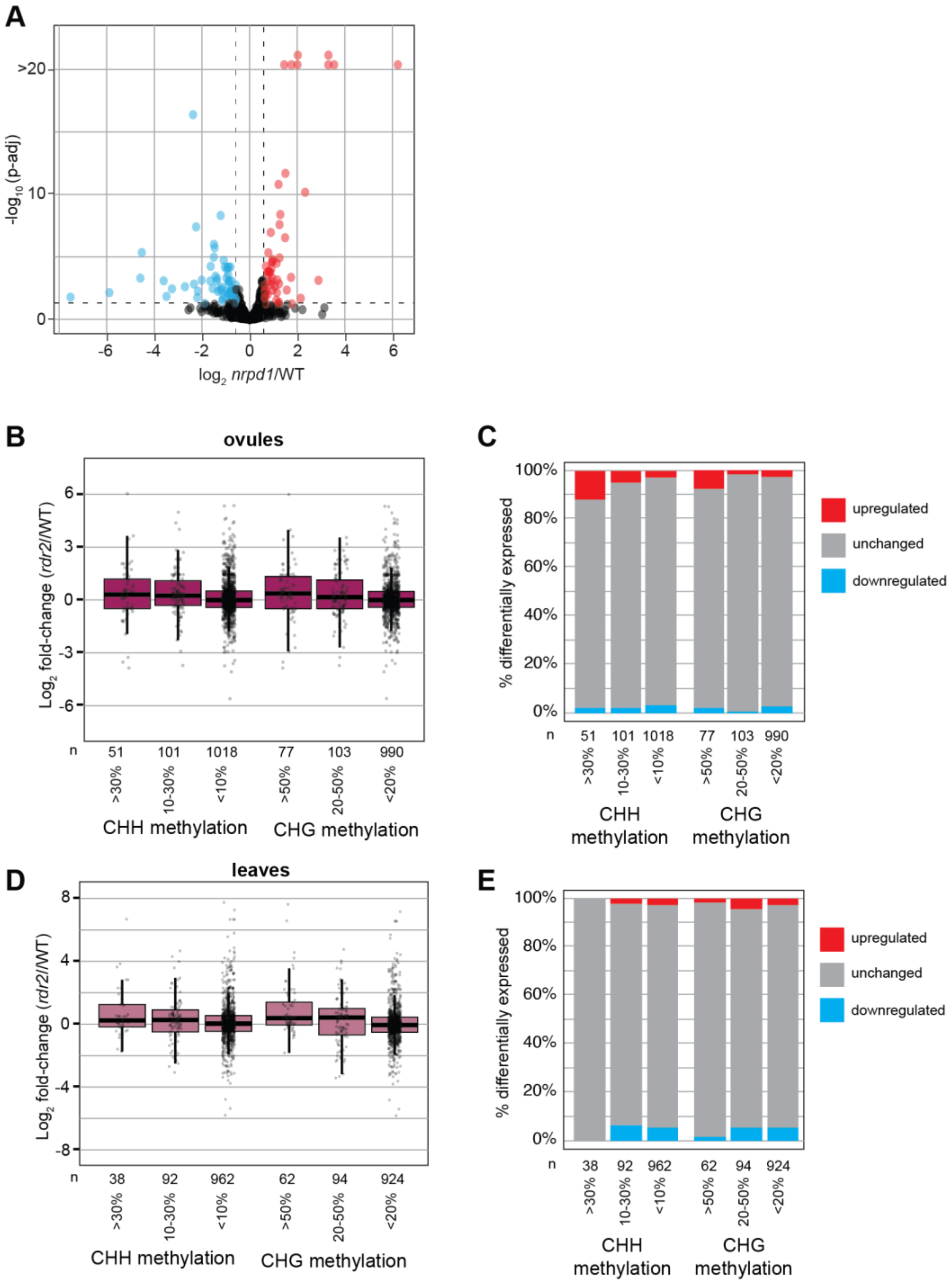


**Supplemental Figure 8. Differential expression in *nrpd1* and *rdr2* ovules and leaves (supporting Figure 5).**

**(A)** Volcano plot of *nrpd1*/WT ovule expression of the 1,270 genes overlapping siren re-aligning windows. Significantly upregulated genes (adjusted p value < 0.05) are shown in red, significantly downregulated genes shown in blue. **(B, D)** $Log_2$ fold-change of *rdr2*/WT ovule (B) or leaf (D) gene expression for the 1,270 genes overlapping siren *trans*-methylation sites, stratified by methylation level. Genes with no detectable expression in either genotype were omitted from the analysis. **(C)** Genes from (B, D) were categorized as upregulated or downregulated when they have an adjusted p-value <0.05. All other genes are considered “unchanged”.

.

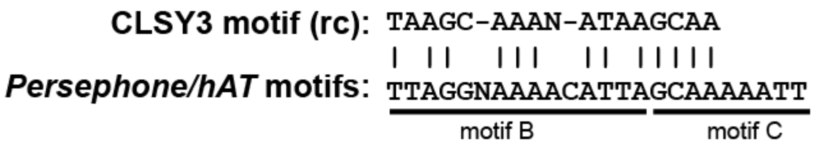


**Supplemental Figure 9. Comparison of siren TE motifs and CLSY3 binding motif (supporting Figure 7).**

The reported CLSY3-associated "motif 1" is similar to motifs B and C that are shared by siren-associated *Persephone* and *Bra_hAT1* elements.

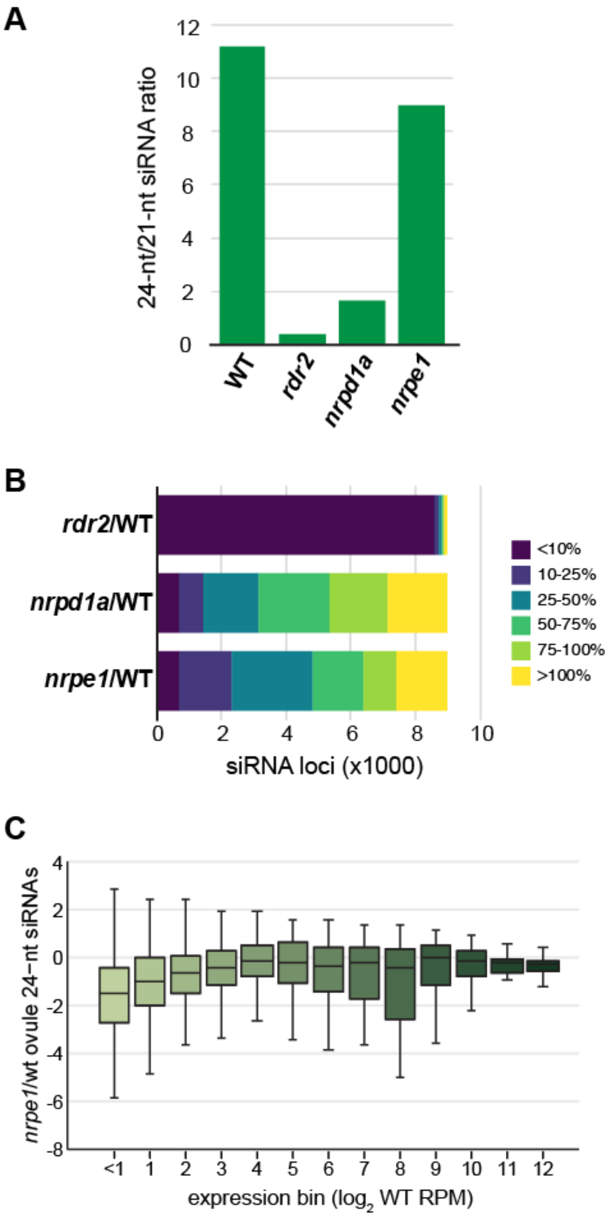


**Supplemental Figure 10. *nrpd1a-2* is required only at the highest-expressing siRNA loci in ovules (supporting Figure 7).**

Although *nprd1a-1* has a reduction in 24-nt siRNA accumulation comparable to *rdr2-2*, most siRNA loci show only a small reduction in siRNA accumulation. **(A)** The ratio of 24-nt to 21-nt siRNAs in *B. rapa* ovules from WT and three RdDM mutants. **(B)** The percent of siRNA expression remaining in ovules of three *B. rapa* RdDM mutants. **(C)** There is no correlation between expression level in wild type and the loss of siRNAs in *nrpe1*. This stands in contrast to *nrpd1a*, for which loci with the highest expression in wild type have the strongest loss of siRNAs (Figure 7C).
